# Positive olfactory childhood memory is rooted in the olfactory bulb and triggers large scale changes beyond the olfactory system

**DOI:** 10.64898/2026.03.19.712996

**Authors:** Jules Dejou, Anna Athanassi, Théo Brunel, Marc Thevenet, Anne Didier, Nathalie Mandairon

**Affiliations:** CNRS, UMR 5292, INSERM, U1028, Lyon Neuroscience Research Center, Neuroplasticity and Neuropathology of Olfactory Perception Team, University of Lyon, 69000; Institut Universitaire de France, Paris, France

**Keywords:** Olfaction, olfactory memory, mice, childhood, neurogenesis, optogenetics, cellular mapping

## Abstract

Olfactory childhood memories are particularly important for forming one’s identity. However, we don’t know how they exert their privileged influence and shape brain structure. To address this, we modeled childhood olfactory memory in mice based on a human survey indicating that our earliest olfactory memory arises from repeated positive experiences paired with a pleasant odorant. Accordingly, mice were exposed during childhood to a pleasant odorant in a playful environment. In adulthood, memory recall relied on neonatal-born granule cells in the olfactory bulb, as their optogenetic silencing impaired retrieval, and on increased functional connectivity in the reward system. With age, memory persistence depended on re-exposure to the childhood odorant and was associated with the disengagement of neonatal-born granule cells, alongside with strengthened limbic functional connectivity. Together, these findings identify neonatal neurons as a key substrate for encoding childhood olfactory memory and reveal dynamic reorganization of brain networks supporting its long-term significance.

## Introduction

Marcel Proust, in his book Swann’s Way (1919), describes the smell of a madeleine dipped in linden tea as triggering intense joy and memories of childhood. This episode clearly illustrates the importance of olfactory childhood memory in the formation of one’s identity. Research in humans has identified two main characteristics of odor-evoked autobiographical memory. First, it tends to originate from an early period of life, typically within the first decade, whereas memories evoked by other sensory modalities, such as visual or auditory cues, are mainly located in the second and third decades of life (Chu & Downes, 2000; Goddard et al., 2005; Hackländer et al., 2019; Miles & Berntsen, 2011; Willander & Larsson, 2007). Second, childhood olfactory memory is generally associated with a more positive emotional valence compared to those triggered by other senses (Arshamian et al., 2013; de Bruijn & Bender, 2018; Hackländer et al., 2019; Herz et al., 2004; Willander & Larsson, 2007).

In humans, the few studies investigating the neural bases of odor-evoked autobiographical memory have shown that this memory elicits greater activation in structures of the limbic system (i.e., amygdala, temporal pole, insula, occipital gyrus) and in regions involved in emotional regulation (i.e., right dorsolateral prefrontal cortex, precuneus, orbitofrontal cortex) compared to exposure to irrelevant odorants, or memories evoked by images and words (Arshamian et al., 2013; Herz et al., 2004). These neuroimaging data coincide with the stronger emotional component of odor-evoked autobiographical memory, compared to memories evoked by other sensory modalities. However, the specific neural mechanisms that govern the formation and long-term persistence of this childhood olfactory memory remain poorly understood.

To investigate its underlying neural bases, we first developed a mouse model of childhood olfactory memory inspired by human data showing that such memory is highly pleasant (Hackländer et al., 2019; Saive et al., 2014; Willander & Larsson, 2007). Conducting our own human online survey, we further complemented these data and showed that childhood olfactory memory relies on repeated – and not single – events and is associated with a pleasant odorant that tends to be re-encountered throughout life. Based on these findings, we modeled a positive olfactory memory in young mice by associating an attractive odorant with an enriched environment between P23 and P33, corresponding to the childhood period. Environmental enrichment has previously been shown to reduce anxiety and promote positive affective states in rodents (Bailoo et al., 2018; Benaroya-Milshtein et al., 2004; Burgess et al., 2013), making it a suitable paradigm to study long-term positive olfactory memory. Using this approach, we first validated the mouse model by showing that the positive environment during childhood was associated with an increased number and frequency of ultrasonic vocalizations, reflecting a positive affective state, and that young adult mice exhibited a preference for the childhood odorant.

In both humans and rodents, olfactory input is first processed in the brain by the olfactory bulb (OB) where the information is finely tuned by local interneurons, the granule cells (GCs) and periglomerular cells (Nagayama et al., 2014). These neurons have the particularity to be mainly formed postnatally: during early infancy in humans and throughout life in rodents, in which proliferation peaks at postnatal day 1 (P1) (Lemasson, 2005). At this stage, progenitor cells located in the OB and in the subventricular zone of the lateral ventricles give rise to neuroblasts that migrate via the rostral migratory stream to the OB, where they differentiate and integrate a developing neuronal network, and show a very high survival rate (Alvarez-Buylla & Garcıia-Verdugo, 2002; Luskin, 1993; Winner et al., 2002). The production of GCs born at P1 can be modulated by early life olfactory experience (Lemasson, 2005; So et al., 2008), and these cells are required for some olfactory learning in adulthood (Forest, Chalençon, et al., 2019; Imayoshi et al., 2008; Sakamoto et al., 2014). This indicates the ability of these neonatal neurons to support the memory traces of olfactory experiences. Importantly, given their enhanced excitability during childhood and high survival rate, we hypothesized that neonatal GCs would be suitable substrates for life-long olfactory memory (Gao & Strowbridge, 2009; Imayoshi et al., 2008; Lemasson, 2005; Sakamoto et al., 2014).

From the OB, the olfactory output is then sent to higher-order olfactory structures, namely the piriform cortex, the anterior olfactory nucleus, the entorhinal cortex, the olfactory tubercle and the cortical amygdala (Hanson et al., 2020; Igarashi et al., 2012; Nagayama, 2010). These structures are closely interconnected with what can be referred to as the olfactory-limbic system, comprising the aforementioned regions along with the accessory olfactory bulb, the basolateral amygdala, the insula and the tenia tecta (Cleland & Linster, 2019; Lane et al., 2020). This olfactory-limbic system is also closely connected to brain regions involved in reward and memory processing, respectively the olfactory tubercle, considered part of the reward circuit (Wesson, 2020; Xiong & Wesson, 2016) and the lateral entorhinal cortex which in turn sends dense projections to the hippocampus (Przy et al., 2025). The significant connections or even overlap between the chemosensory circuit and the brain regions involved in value processing and memory may explain the affective response elicited by these odorants.

Finally, we investigated the retention of the olfactory memory in later adulthood and the associated network reorganization.

Our results revealed that P1-born GCs coded the olfactory memory, as indicated by their enhanced recruitment in response to the childhood odorant, while their photo-inhibition altered the preference for this odorant in adults. We then analyzed in young adults the cFos expression across 27 brain regions involved in olfactory-limbic, reward and memory circuits. We revealed that correlated activities increased within and between the reward and memory systems in response to the childhood odorant, in comparison to the control condition, likely contributing to the unique vividness and strong affective potency of childhood olfactory memory. In later adulthood, the persistence of the memory required periodic re-exposures to the childhood odorant but no longer preferentially recruited P1-born GCs. The functional brain connectivity was reshaped with enhanced connectivity within the olfactory-limbic system along with the medial prefrontal cortex, the orbitofrontal cortex as observed in younger adults, but with a disengagement of the dorsal hippocampus.

## Results

### Modeling childhood olfactory memory in mice: evidence from a human survey

To model a childhood olfactory memory in mice, we first conducted an online survey in humans to better define its features. We asked 647 participants to recall an odorant that was significant in their childhood, going back as far as possible in time. To evaluate the emotional valence of the initial event, we asked participants to rate the pleasantness and the six basic emotions (happiness, surprise, fear, disgust, sadness, anger) on 1-9 scales (Ekman, 1992). The ratings differed between emotions (F(2.51, 1618.67) = 920.87, p < 0.0001) and were the highest for positive emotions namely happiness and pleasantness, followed by surprise and then negative emotions namely fear, disgust, sadness and anger (**Fig. 1A**). Then, to estimate the number of exposures to the initial context required to form the childhood olfactory memory in mice, participants were asked how many times the event that led to their earliest olfactory memory had occurred. 73.1% of participants reported that these events had occurred more than five times, a proportion of subjects significantly higher than those reporting events repeated between 2 to 5 times (13.4%, p < 0.0001) and single events (13.4%, p < 0.0001; **Fig. 1B**). Next, to guide the selection of odorants for the mouse model, participants had to rate the hedonic value of their childhood odorant, which was found to be predominantly pleasant (mean = 7.1 on a 1-9 scale; **Fig. 1C**). We then investigated whether the hedonic ratings of the odorants reflected their intrinsic pleasantness rather than being retrospectively influenced by a positive emotional context. For that purpose, we used the dataset from (Dravnieks et al., 1984) which provides estimated hedonic values for 146 odor descriptors, 51 of which were reported by our participants. We observed that the estimated hedonic values of the descriptors identified in our survey were significantly higher than those of the descriptors not reported (one-tailed t-test, t(98.9) = 2.01, p = 0.024, d = 0.35;).

**Fig. 1.**
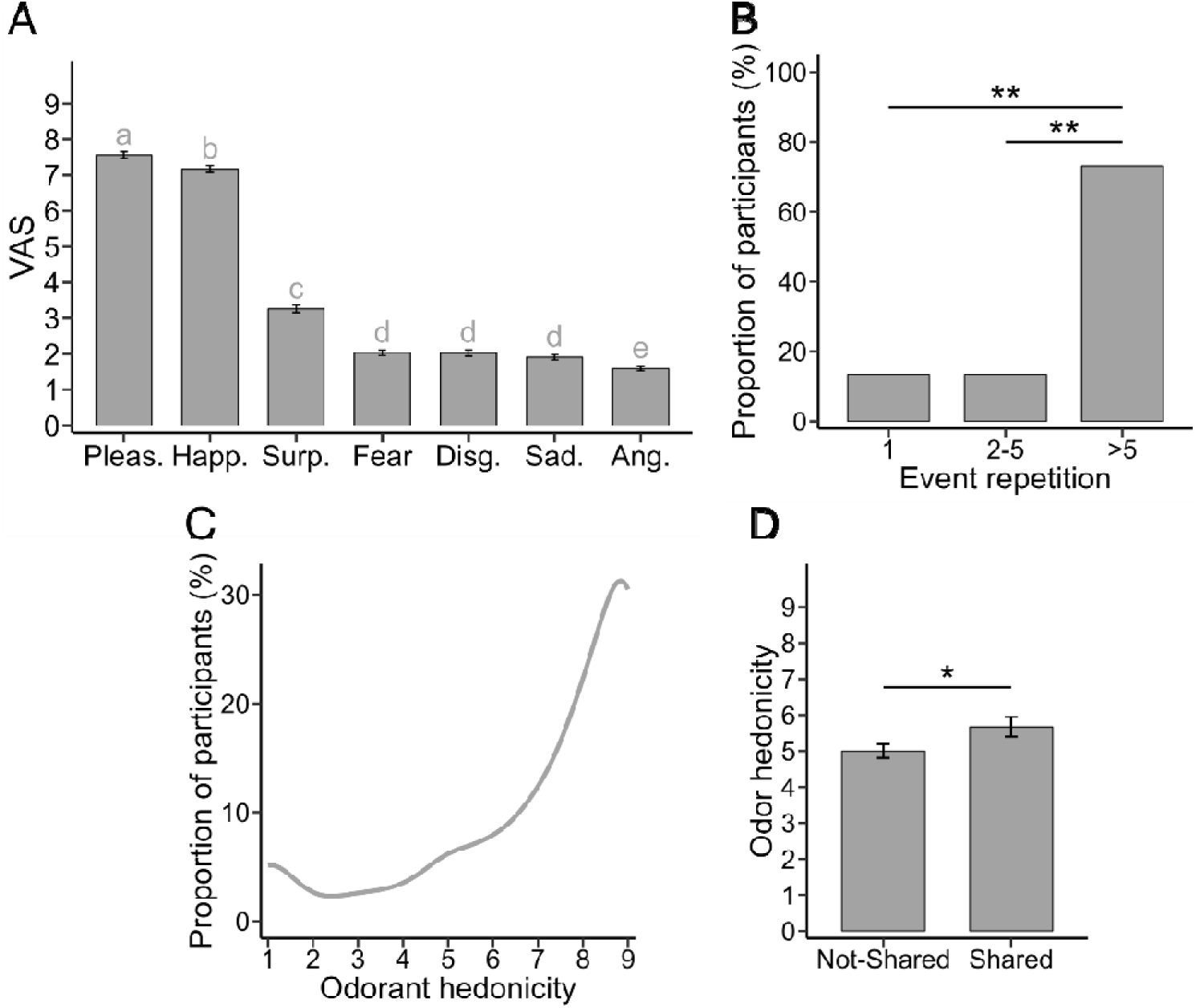
Childhood olfactory memory arises from repeated positive events associated with a spontaneously pleasant odorant. The initial event of the childhood olfactory memory is (**A**) associated with greater positive emotions compared to negative ones, and is (**B**) reported to have occurred more than five times by most participants. (**C** and **D**) Hedonic value of the childhood odorant. (**C**) The hedonic value of the childhood odorant is highly positive. (**D**) Panelists rate the hedonic value of odorants present in our survey (n=51) as more pleasant than those absent from it (n=95). Data are represented as data points and mean ± SEM. Statistical significance depicted as **p < 0.01. Letters represent statistical groups resulting from the post-hoc analysis. Pleas. = Pleasantness, Happ. = Happiness, Surp. = Surprise, Disg. = Disgust, Sad. = Sadness, Ang. = Anger.

This supports the notion of odor–memory congruency, whereby individuals preferentially encode pleasant odorants into autobiographical memory. Based on these findings, to install a long-term childhood memory, mice were exposed from P23 to P33, once to a positive environment without a paired odorant, followed by five exposures to the same context paired with an odorant known as attractive for them (either (+)-limonene, β-citronellol or camphor) (Chalençon et al., 2024; Kermen et al., 2016; Midroit et al., 2021).

### Olfactory memory is built in a positive affective context

To establish a childhood olfactory memory, P23 mice were exposed to a pleasant odorant in an enriched environment which served as the positive context. More precisely, they were placed in a large, enriched cage designed to be playful and promote exploration and social interactions (PLAY-O condition). A control group was exposed to the same odorant in regular housing condition (CTRL-O condition). These exposures were done 5 times (every other day from P23 to P33) each lasting two hours (**Fig. 2A**). A way to assess the emotional state in rodents is to record their ultrasonic vocalizations (USVs). Indeed, it has been demonstrated that positive affective states induce longer, higher-frequency and higher-number USVs in rodents compared to negative affective states (Granon et al., 2018; Kuwaki & Kanno, 2021; Lefebvre et al., 2020). We thus recorded the USVs of PLAY-O and CTRL-O mice in their respective enriched and control housing environment, and revealed that the number (one-tailed t-test, t(15.53) = -2.42, p = 0.014; **Fig. 2B**) and frequency (one-tailed t-test, t(16) = -1.98, p = 0.03; **Fig. 2C**) of USVs emitted in the experimental setup were higher in the PLAY-O group compared to the CTRL-O group. No difference in USVs duration was observed (one-tailed t-test, t(9.56) = 0.22, p = 0.58; **Fig. 2D**). These results suggest that mice experienced a more positive affective state in the enriched environment compared to the standard housing condition.

**Fig. 2.**
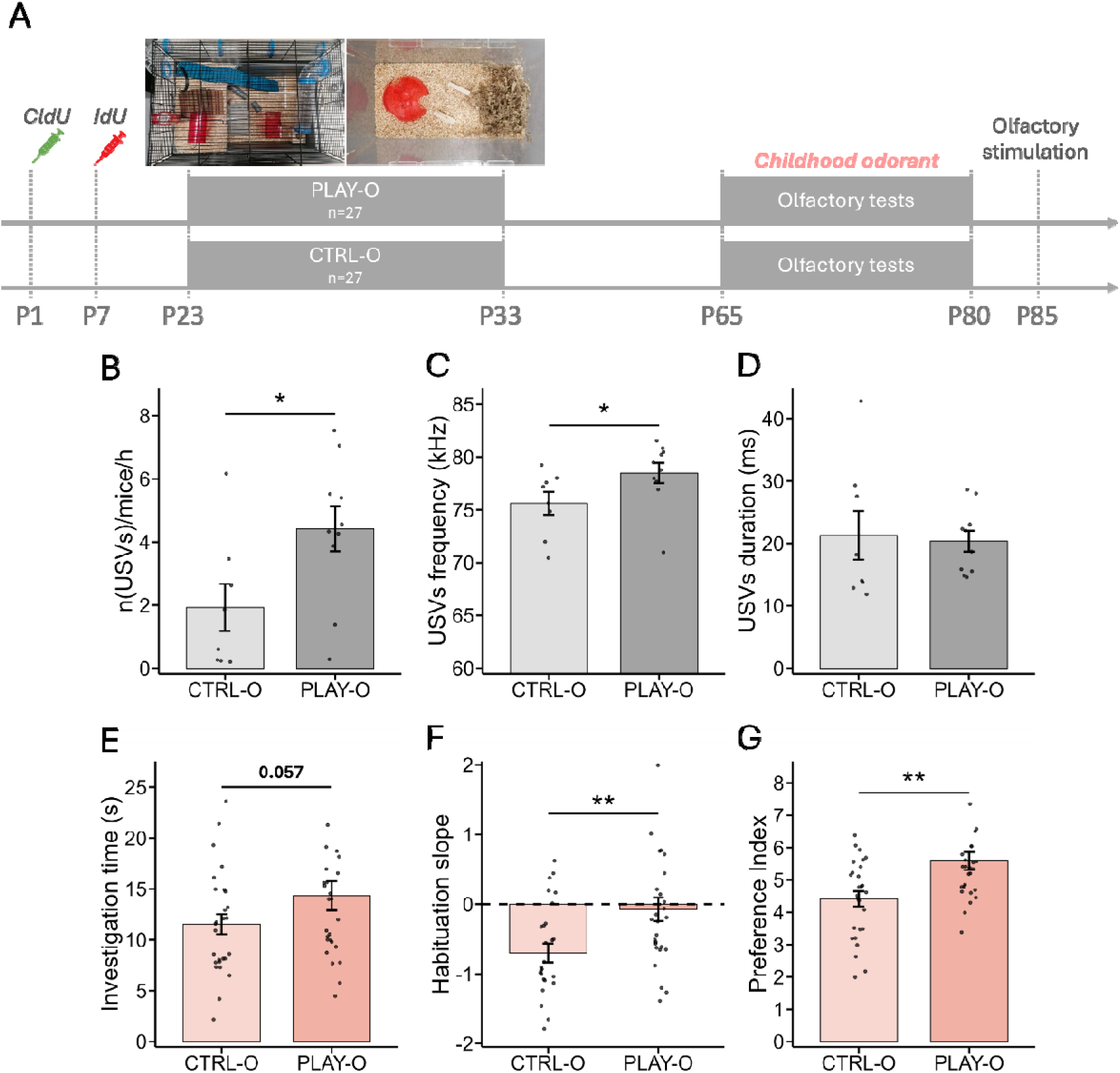
The childhood odorant is encoded in a positive context and is preferred in adulthood. (**A**) PLAY (upper left) and CTRL (upper right) setups. Timeline of the experiment (lower). (**B** and **D**) Multi-individual recording of ultrasonic vocalizations. PLAY-O (n=10 cages) mice show an increase in (**B**) the number (**C**) the frequency, but not (**D**) the duration of the USVs compared to CTRL-O mice (n=8 cages). (**E** to **G**) Behavioral responses to the childhood odorant. (**E**) PLAY-O mice investigate more the odorized hole containing the childhood odorant in the exploration task compared to CTRL-O mice. (**F**) The slope of odorant habituation is higher for PLAY-O (n=27) compared to CTRL-O (n=27) mice. (**G**) PLAY-O mice show an increased preference index for the childhood odorant compared to CTRL-O mice. Data are represented as data points and mean ± SEM. Statistical significance depicted as *p < 0.05, **p□<□0.01.

### The childhood odorant is remembered in adulthood with a positive affective valence

Olfactory preference for the childhood odorant was assessed in adulthood using two behavioral tests. First, the odorant exploration test was used to evaluate, in a single 2-minute trial, childhood odorant attractiveness (Chalençon et al., 2024; Kermen et al., 2016; Mandairon et al., 2009; Midroit et al., 2021). Second, we made the hypothesis that the childhood odorant would be preferred to the point of counteracting the loss of interest to the odorant classically observed across repeated exposures. We then used the habituation test which consists in presenting the childhood odorant across four consecutive 50-second trials to evaluate changes in odorant investigation time – typically decreasing as the odorant becomes familiar. The exploration test revealed that the exploration time of the childhood odorant tended to be higher in the PLAY-O group compared to the CTRL-O group (one-tailed t-test, t(52) = -1.610, p = 0.057; **Fig. 2E**), reflecting a preference for the childhood odorant. Analysis of the habituation slope revealed a significantly higher habituation coefficient in PLAY-O compared to CTRL-O mice (one-tailed t-test, t(52) = -2.899, p = 0.0027; **Fig. 2F**) indicating a lack of habituation in the PLAY-O group, likely reflecting increased preference of the odorant. These results, combined in a preference index (i.e., sum of z-scored values from both tests) showed an increased preference for the childhood odorant of the PLAY-O compared to the CTRL-O condition (one-tailed Wilcoxon test, W = 210, p = 0.0035; **Fig. 2G**). These data indicate that PLAY-O group showed a positive memory of the childhood odorant. Next, to investigate whether the preference observed in adulthood was specific to the childhood odorant, we tested the preference of PLAY-O and CTRL-O mice in response to an unknown odorant. We found that the exploration time of the odorant in the exploration test was similar between PLAY-O and CTRL-O groups (two-tailed t-test, t(43.57) = -1.369, p = 0.169; **Fig. S1A**). In addition, no difference was found regarding the habituation slope (two-tailed Wilcoxon test, W = 299.5, p = 0.265; **Fig. S1B**). In line with these results, the preference index resulting from the two behavioral tests was similar between both groups (two-tailed Wilcoxon test, W = 261, p = 0.077; **Fig. S1C**). These data indicate that the preference observed for PLAY-O mice is specific to the childhood odorant. To further ensure that the observed preference for the childhood odorant was not solely driven by the enriched environment itself, but rather by its association with the odorant, we added two experimental groups: one exposed to the enriched environment and one maintained in the regular housing, both without an associated odorant (PLAY-NO and CTRL-NO groups, **Fig. S2A**). In response to an odorant (that is unknown), we found similar odorant investigation time in the exploration test (two-tailed Wilcoxon test, W = 409.5, p = 0.139) and habituation slope (two-tailed t-test, t(63) = 1.270, p = 0.219) between PLAY-NO and CTRL-NO groups, leading to similar preference index (two-tailed t-test, t(63) = -0.041, p = 0.968) (**Fig. S2B-D**).

### The childhood olfactory memory enhances the functional recruitment of P1-born granule cells in the olfactory bulb of young adult mice

In the OB, GCs are formed mainly postnatally with a peak of intense proliferation at P1 (Bayer, 1983; Lemasson, 2005), making them maturing in the OB concomitantly to olfactory childhood experiences. To explore the role of P1-born GCs in the childhood olfactory memory, we labeled them using CldU injection and we analyzed their cFos expression in response to the childhood odorant. We revealed that cFos expression of P1-born GCs (CldU-positive cells) was higher in response to the childhood odorant in the PLAY-O compared to the CTRL-O group (one-tailed t-test, t(30) = -2.326, p = 0.014; **Fig. 3A**). Importantly, no difference in cFos expression in P1-born granule cells was observed between PLAY-NO and CTRL-NO groups in response to an unknown odorant (two-tailed t-test, t(18) = 0.958, p = 0.351; **Fig. S3**), confirming the preferential recruitment of P1-born neurons in response to the childhood odorant. This enhanced functional recruitment of P1-born GCs was not due to a generalized increase in cellular activity in the OB since cFos-positive cell density in the granule cell layer was not different between PLAY-O and CTRL-O groups (two-tailed t-test, t(30) = 0.293, p = 0.772; **Fig. 3B**). Finally, the CldU-positive cell density was similar between PLAY-O and CTRL-O mice, indicating no effect of the childhood olfactory memorization on the survival of P1-born GCs (two-tailed t-test, t(30) = -0.012, p = 0.99; **Fig. 3C**).

**Fig. 3.**
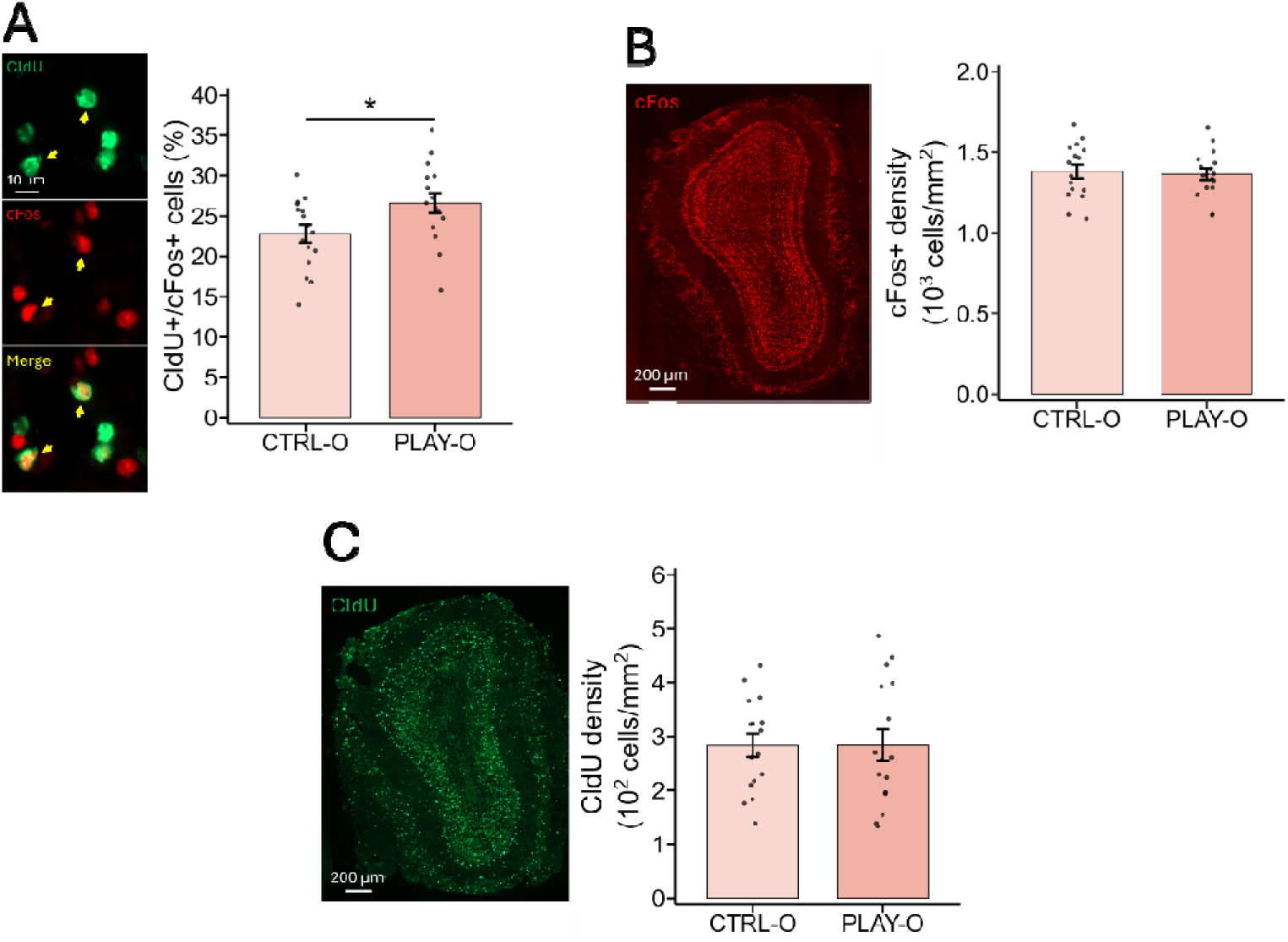
The childhood olfactory memory functionally recruits P1-born GCs of the OB. (**A**) Example of activated P1-born GCs (CldU/cFos-positive cells: yellow arrows, upper). The percentage of CldU-positive cells responding to the childhood odorant is higher for PLAY-O (n=16) compared to CTRL-O (n=16) mice (lower). (**B**) Example of a section with cFos labeling of the GCL (left). The density of cFos-positive cells in the GCL of the OB is similar between PLAY-O (n=16) and CTRL-O (n=16) mice (right). (**C**) Example of a section with CldU labeling of the GCL (left). The density of CldU-positive cells in the GCL of the OB is similar between PLAY-O (n=16) and CTRL-O (n=16) mice (right). Data are represented as data points and mean ± SEM. Statistical significance depicted as *p□<□0.05.

### The optogenetic inhibition of P1-born granule cells prevents the recall of childhood olfactory memory

To explore the functional role of P1-born GCs in the childhood olfactory memory, we inhibited them using optogenetic tools. For that purpose, two new groups of mice received an intraventricular injection of a lentivirus expressing the inhibitory photo-sensitive channel halorhodopsin (PLAY-Halo) or a control virus (PLAY-Cont) at P1, to transduce P1-born GCs. The two groups were exposed to an odorant in a positive, enriched environment during childhood (PLAY-O condition) as previously described, and they were light-stimulated in adulthood during the behavioral tasks to assess the impact of P1-born GCs inhibition on odorant preference (**Fig. 4A**). The photo-inhibition applied each time the mice explored the childhood odorant resulted in a decrease in the odorant investigation time in the exploration test (one-tailed t-test, t(28) = 1.94, p = 0.031; **Fig. 4B**) and a reduced habituation slope in the habituation test (one-tailed Wilcoxon test, W = 149, p = 0.043; **Fig. 4C**) for the PLAY-Halo (inhibited) compared to the PLAY-Cont (non-inhibited) group. Consequently, the preference index decreased in the PLAY-Halo compared to the PLAY-Cont group (one-tailed t-test, t(28) = 1.95, p = 0.03; **Fig. 4D**). This reduction in preference was specific to the childhood odorant as it was not observed for an unknown odorant (exploration test: two-tailed t-test, t(28) = 0.58, p = 0.57; habituation test: two-tailed t-test, t(28) = -0.23, p = 0.82; preference index: two-tailed t-test, t(28) = 0.21, p = 0.83; **Fig. S4**). Then, we assessed the level of viral transfection by analyzing the density of EYFP-positive cells in the granule cell layer of the OB and found no difference between PLAY-Halo and PLAY-Cont groups (two-tailed t-test, t(16) = 1.16, p = 0.26; **Fig. 4E-F**). We also analyzed the effectiveness of light-induced inhibition of P1-born GCs and showed a decrease in EYFP/cFos-positive cell density in the granule cell layer for the PLAY-Halo compared to the PLAY-Cont group (one-tailed t-test, t(16) = 2.40, p = 0.01; **Fig. 4G**). Together, these data indicate that the inhibition of P1-born GCs prevents the childhood olfactory memory recall and highlights their prominent role in the encoding and recall of the olfactory memory.

**Fig. 4.**
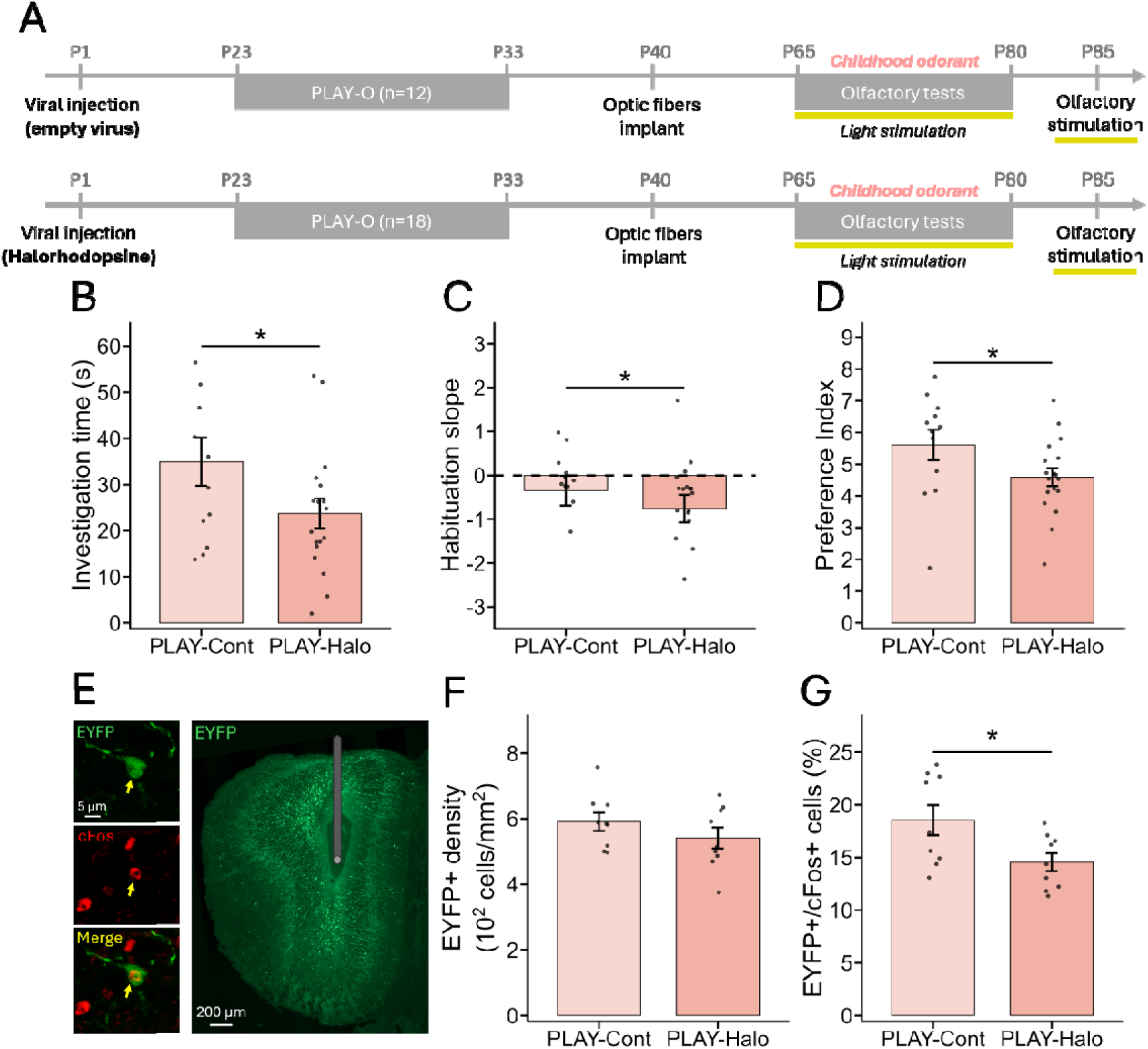
Optogenetic inhibition of P1-born GCs prevents the preference for the childhood odorant. (**A**) Experimental timeline of the optogenetic experiment. (**B** to **D**) Behavioral responses to the childhood odorant. The light-induced inhibition decreases (**B**) the investigation time, (**C**) the habituation slope as well as (**D**) the preference index for PLAY-Halo (n=18) compared to PLAY-Cont (n=12) mice in response to the childhood odorant. (**E** to **G**) Transduced P1-born GCs activity and density. (**E**) Example of an EYFP/cFos-positive cell (yellow arrow, left). Example of a section with EYFP labeling of P1-born GCs, with a schematic representation of the optic fiber location (right). (**F**) The density of EYFP-positive cells is similar between PLAY-Cont (n=9) and PLAY-Halo (n=9) groups. (**G**) The light stimulation significantly decreases the percentage of EYFP/cFos-positive cells in PLAY-Halo compared to PLAY-Cont mice. Data are represented as data points and mean ± SEM. Statistical significance depicted as *p < 0.05.

### The childhood odorant enhances the cerebral functional connectivity in young adult mice

The OB projects to several brain structures that are monosynaptically or indirectly connected to regions belonging to the olfactory-limbic, memory, reward systems and are thought to be involved in olfactory memory (Cleland & Linster, 2019; Kontaris et al., 2020; Soudry et al., 2011), as well as cortical areas. To better characterize the neural bases of childhood olfactory memory, we analyzed brain activation and functional connectivity through cFos-positive cell mapping, to identify network remodeling associated with the childhood olfactory memory (**Fig. 5A**). We therefore selected a total of 27 regions, grouped into 4 functional systems named “olfactory-limbic”, “memory”, “reward” and “cortical areas” (see Material and Method section for a detailed list of the structures). First, after thresholding and assigning cFos-labeled cells to the selected brain regions, multiple Wilcoxon tests performed on each region revealed no significant differences in cFos-positive cell density between PLAY-O and CTRL-O groups (**Fig. 5B**, **Table S1**). This suggests that the observed preference for the childhood odorant did not change the mean level of activity in these regions. Then, we examined how activity co-varied across the 27 regions in each of the two experimental groups considering that co-variation reflects functional connection. Correlations in cFos-positive cell density were calculated for all region pairs within each group, generating corresponding correlation matrices for PLAY-O and CTRL-O groups (**Fig. S5A-B**). We also computed comparison matrices (PLAY-O – CTRL-O) to visualize the differences in correlation strength (**Fig. S5C**). A significance threshold of p < 0.01 was then applied to retain only robust correlations. The corresponding network graphs were generated for the PLAY-O and CTRL-O groups (**Fig. 5C-D**), and comparison matrices were constructed to visualize group-specific connections (**Fig. S5D**). Functional connectivity analyses revealed that exposure to the childhood odorant induced a distinct functional organization in the PLAY-O group compared to the CTRL-O group, characterized by an enhanced recruitment of the memory system along with its interaction with the reward system. In fact, calculation of the total connection density for each group (i.e., the number of connections realized over the total possible) revealed an increase in the memory (p = 0.0009) and reward (p = 0.038) systems for PLAY-O compared to CTRL-O group (**Fig. 5E**). Specifically, the increase in connectivity in the memory system was partly driven by enhanced intra-system connection density (p = 0.023, **Fig. 5F**), whereas for the reward system, this was supported by increased inter-system connections with the memory system (p = 0.017, **Fig. 5G**). The other comparisons were not significant (**Table S2**). To investigate the weight of each system in the established network, we analyzed the relative connection density (i.e., the number of connections normalized by total connections observed in each group). The results are consistent with the previous ones, highlighting a greater influence of the memory system (overall, p = 0.0076; intra-memory, p = 0.054) and its interactions with the reward system (p = 0.074). They also show an increased contribution of the olfactory-limbic system in the CTRL-O compared to the PLAY-O group (overall, p = 0.014; intra-olfactory-limbic, p = 0.069; olfactory-limbic-cortex, p = 0.081) (**Fig. S5E-G**). We confirmed the robustness of these results (i.e., total and relative connection densities) by repeating the analyses using two different thresholds (p < 0.005 and p < 0.025) (**Table S2**). To get insights into how the different brain regions interact as a system, we generated PLAY-O and CTRL-O networks using Gephi software, enabling the representation of the Louvain community subnetworks (i.e., method that identifies groups of nodes that are more strongly connected to each other than to the rest of the network). Qualitative analysis of the networks interestingly showed that the PLAY-O group had a subnetwork composed of the MOB, dHipp, OFC and CPu, which was directly connected to the mPFC via the MOB and dHipp (**Fig. 5H-I**). To confirm that the observed functional connectivity changes were specific to the childhood olfactory memory and not to the environmental enrichment, PLAY-O mice were compared to PLAY-NO mice. cFos-positive cell density analysis revealed higher activity in the dHipp (Wilcoxon test, p = 0.041) and S1 (Wilcoxon test, p = 0.045) in PLAY-O compared to PLAY-NO mice (**Fig. S6A**, **Table S3**). The analysis of the total and relative connection densities revealed an increased connectivity in the memory and reward systems, mainly through their interaction, while the connectivity was decreased in the olfactory-limbic system for PLAY-O compared to PLAY-NO mice (**Fig. S6B-G**, **Table S4**). The Gephi representation showed that the PLAY-O-specific subnetwork involving MOB, dHipp, OFC, CPu and mPFC was absent in PLAY-NO mice (**Fig. S6H-I**). Altogether, the findings indicate that the childhood olfactory memory is associated with a strong involvement of memory and reward systems. Interestingly, we observed a well-known subnetwork composed of structures typically involved in autobiographical memory (dHipp-mPFC-OFC) (Reagh & Ranganath, 2018; Saive et al., 2014; Sekeres et al., 2018), also including the CPu and MOB which shows the involvement of olfactory modality and may explain part of the emotional potency of childhood olfactory memory.

**Fig. 5.**
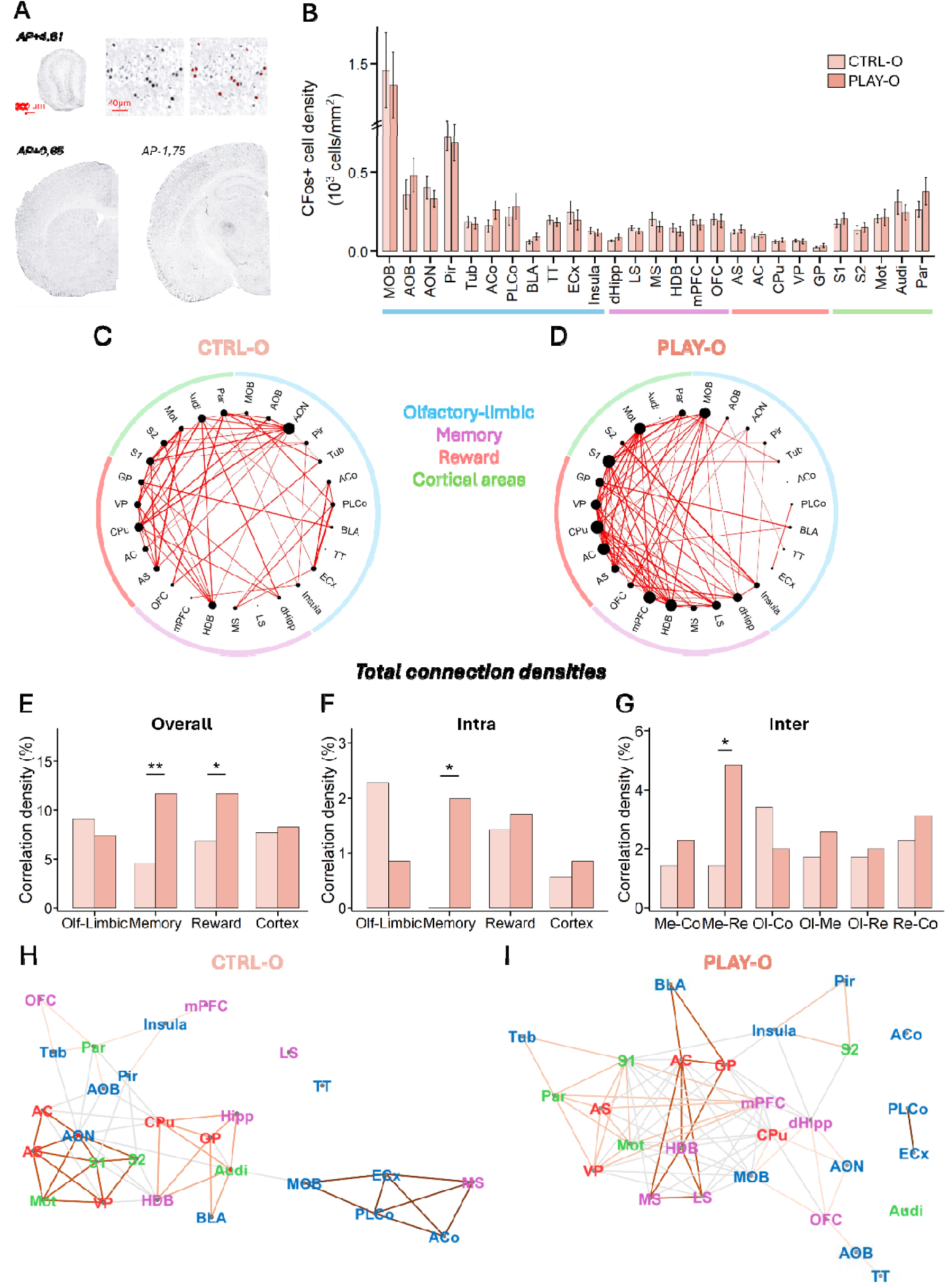
Exposure to the childhood odorant functionally recruits memory and reward systems. (**A**) Example of sections with cFos labeling at anteroposterior coordinates +4.61 (upper left), +0.65 (lower left), -1.75 (lower right). Example of cFos-positive cells labeling and detection (upper right). (**B**) cFos-positive cell density was similar between PLAY-O (n=10) and CTRL-O (n=10) mice in the selected brain regions. (**C-D**) Representation of functional connectivity networks in response to the childhood odorant, constructed by considering correlations with Pearson’s p < 0.01. Each region is represented by a dot; the size of the dot is a function of the number of connections. The thickness of the line increases with the associated correlation coefficient. These networks are represented for (**C**) CTRL-O (n=10) and (**D**) PLAY-O (n=10) groups. (**E-G**) Total correlation density (i.e., the number of connections realized over the total possible) analysis. (**E**) The total correlation density is increased for PLAY-O compared to CTRL-O group in memory and reward systems. This increase in total correlation density is particularly observed in (**F**) intra-memory system as well as (**G**) memory-reward inter-system. (**H-I**) Representation of the Gephi networks with Louvain Communities for (**H**) CTRL-O and (**I**) PLAY-O groups. Text colors represent the systems to which the structures belong. The colored lines represent the subnetworks identified with the Louvain Community approach. Statistical significance depicted as *p < 0.05, **p < 0.01. AC = Accumbens Core; ACo = Anterior Cortical Amygdala; AOB = Accessory Olfactory Bulb; AON = Anterior Olfactory Nucleus; AS = Accumbens Shell; Audi = Auditory Cortex; BLA = Basolateral Amygdala; CPu = Caudate Putamen; dHipp = dorsal Hippocampus; GP = Globus Pallidus; HDB = Horizontal Limb of the Diagonal Band of Broca; LS = Lateral Septum; MOB = Main Olfactory Bulb; Mot = Motor Cortex; mPFC = medial Prefrontal Cortex; MS = Medial Septum; OFC = Orbitofrontal Cortex; Par = Parietal Cortex; ECx = Entorhinal Cortex; Pir = Piriform Cortex; PLCo = Posterolateral Cortical Amygdala; S1 = Somatosensory Cortex 1; S2 = Somatosensory Cortex 2; Tub = Olfactory Tubercle; TT = Tenia Tecta; VP = Ventral Pallidum.

### Olfactory memory fades over time

We next sought to understand how the olfactory memory persists through time. To this end, we used a separate cohort of mice that underwent the same experimental protocol as previously described (PLAY-O and CTRL-O) with the exception that olfactory memory was assessed at 6 months of age (**Fig. 6A**). At this stage, PLAY-O mice no longer exhibited increased preference for the childhood odorant compared to CTRL-O mice. This was evidenced by similar investigation time during the exploration test (one-tailed Wilcoxon test, W = 463, p = 0.202; **Fig. 6B**), similar habituation slopes (one-tailed t-test, t(63) = 0.447, p = 0.672; **Fig. 6C**), and similar preference index in response to the childhood odorant (one-tailed t-test, t(63) = - 0.317, p = 0.376; **Fig. 6D**) between PLAY-O and CTRL-O groups. When tested for an unknown odorant, PLAY-O and CTRL-O groups showed similar results on exploration test (two-tailed Wilcoxon test, W = 329.5, p = 0.343), habituation test (two-tailed t-test, t(63) = -0.067, p = 0.947), and preference index (two-tailed t-test, t(63) = -0.531, p = 0.597) (**Fig. S7A-C**). At the cellular level, PLAY-O and CTRL-O groups showed no difference in the percentage of activated P1-born GCs (CldU/cFos-positive cells, two-tailed Wilcoxon test, W = 36, p = 0.497; **Fig. 6E**) in response to the childhood odorant. Together, these findings indicate that the olfactory memory naturally fades over time, in association with a disengagement of P1-born GCs.

**Fig. 6.**
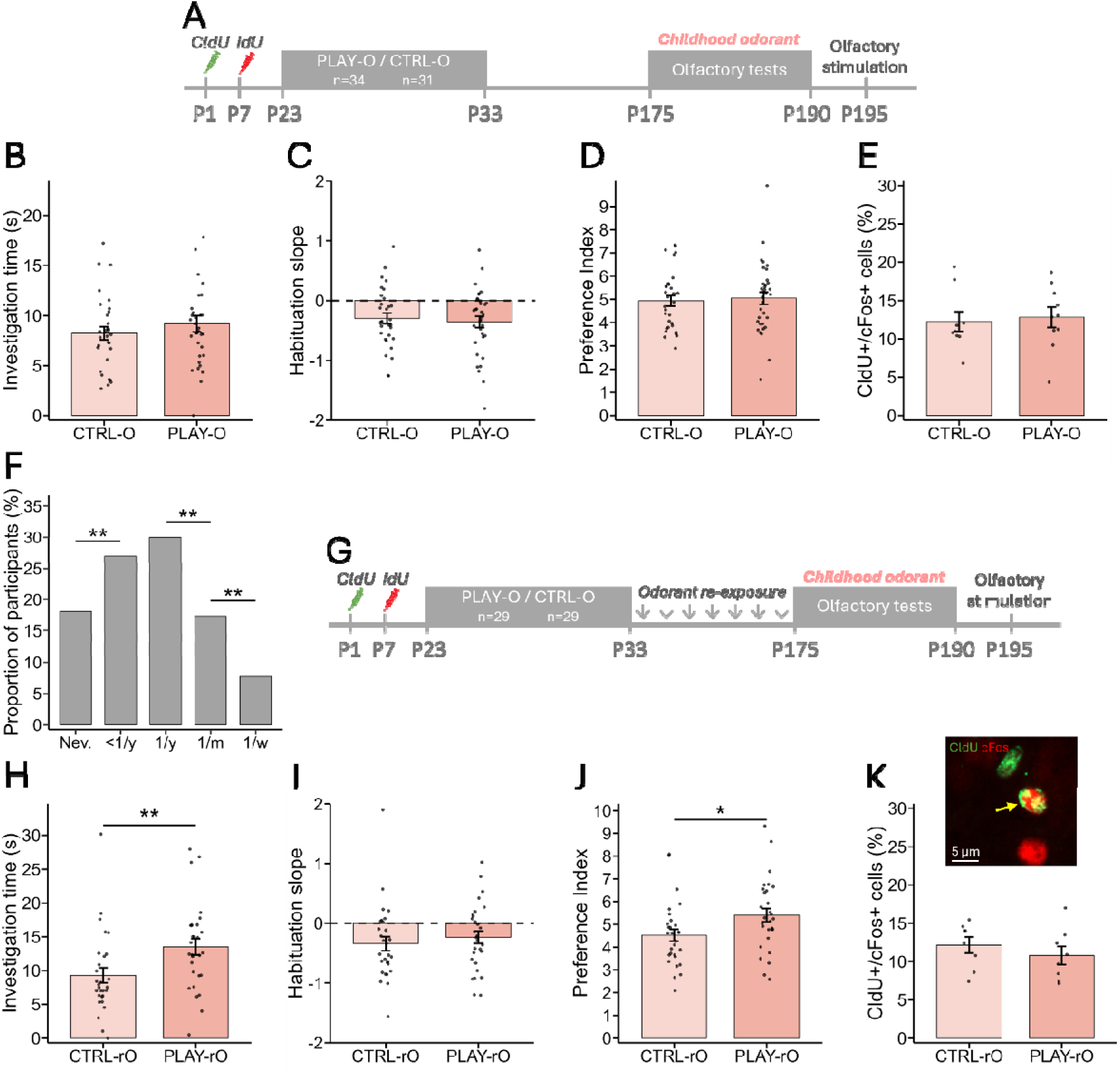
Long-term olfactory memory requires olfactory re-exposures through time and does not rely on P1-born GCs anymore. (**A**) Experimental timeline of the non-exposed groups. (**B-D**) Behavioral responses to the childhood odorant at 6 months of age. PLAY-O (n=34) and CTRL-O (n=31) mice show similar (**B**) investigation time, (**C**) habituation slope and (**D**) preference index regarding the childhood odorant. (**E**) The percentage of CldU-positive cells responding to the childhood odorant shows no difference between PLAY-O (n=10) and CTRL-O (n=9) mice. (**F**) The human survey reveals that the frequency of re-exposure to the childhood odorant is unevenly distributed in the population, with the odorant being periodically re-encountered. (**G**) Experimental timeline of the re-exposed groups. (**H-J**) Behavioral responses to the childhood odorant at 6 months of age, with periodic re-exposures. (**H**) PLAY-rO (n=29) mice display an increased investigation time of the childhood odorant compared to CTRL-rO (n=29) mice. (**I**) The habituation slope is similar between the two groups. (**J**) The resulting preference index is increased for PLAY-rO compared to CTRL-rO mice. (**K**) Example of a CldU/cFos-positive cell (upper). PLAY-rO (n=8) and CTRL-rO (n=8) groups do not show any difference in terms of percentage of CldU/cFos-positive cells (lower). Data are represented as data points and mean ± SEM. Statistical significance depicted as *p < 0.05, **p < 0.01.

### Repeated exposures to the childhood odorant throughout life prevents the decline of olfactory memory

Interestingly, the survey data collected from human participants revealed that the odorant associated with the childhood olfactory memory is periodically re-experienced throughout life by 82% of participants (**Fig. 6F**). Based on these findings, we re-exposed PLAY-O and CTRL-O mice to their childhood odorant in the regular housing every 3 weeks for 2-hour periods, from 2 to 6 months of age (7 re-exposures in total; PLAY-rO and CTRL-rO groups, **Fig. 6G**). These recalls allowed a sustained positive memory for the childhood odorant in PLAY-rO mice compared to CTRL-rO mice, as revealed by an increased investigation time of the childhood odorant in the exploration test (one-tailed Wilcoxon test, W = 240, p = 0.0023; **Fig. 6H**), an increased habituation slope which did not reach statistical significance (one-tailed t-test, t(56) = - 0.693, p = 0.246; **Fig. 6I**), and an increased preference index (one-tailed t-test, t(56) = - 2.239, p = 0.015; **Fig. 6J**) in the PLAY-rO compared to the CTRL-rO group. We confirmed the specificity of the observed increased preference for the childhood odorant, as PLAY-rO and CTRL-rO groups showed similar results on the exploration test (two-tailed t-test, t(56) = 0.067, p = 0.947), habituation test (two-tailed t-test, t(54) = -0.736, p = 0.465), and preference index (two-tailed t-test, t(54) = -0.426, p = 0.671) for an unknown odorant (**Fig. *S*8A-C**). Surprisingly, this sustained memory was not associated with a higher cFos activity of P1-born GCs (one-tailed t-test, t(14) = 0.851, p = 0.795, **Fig. 6K**). Together, the present findings suggest that the childhood olfactory memory has the ability to persist over time through periodic re-exposures to the childhood odorant, but is accompanied by a disengagement of P1-born GCs of the OB.

### The preference for the childhood odorant in later adulthood is associated with a brain functional reorganization recruiting the olfactory-limbic system

To investigate how the functional organization of the brain in response to the childhood odorant evolves with time, we analyzed brain activity and functional connectivity within the same regions of interest belonging to olfactory-limbic, memory and reward systems as well as cortical areas, as previously described. First, cFos-positive cell density was assessed for the selected brain regions in the two experimental groups re-exposed to the childhood odorant throughout life (PLAY-rO and CTRL-rO). Multiple Wilcoxon tests conducted on each selected brain region revealed no significant differences in cFos-positive cell density between PLAY-rO and CTRL-rO groups (**Fig. 7A**, **Table S5**). Next, we investigated how functional connectivity was modulated in response to the childhood odorant by computing correlation matrices and network graphs as previously described (**Fig. 7B-C**, **Fig. S9A-D**). Functional connectivity analyses on thresholded data (p < 0.01) revealed that exposure to the childhood odorant induced a distinct functional organization in the PLAY-rO group compared to the CTRL-rO group, characterized by an enhanced recruitment of the olfactory-limbic system. In fact, calculation of both total connection density (i.e., the number of connections realized over the total possible) and relative connection density (i.e., the number of connections normalized by total connections observed in each group, to assess the weight of each system in the network) for each group revealed an increased connectivity for the olfactory-limbic system in the PLAY-rO group. This increase encompassed both intra-system connections and interactions with cortical areas. The connectivity was decreased for memory and reward systems as well as the interaction between the reward system and cortical areas (**Fig. 7D**-**F**, **Fig. S9E-G**, **Table S6**). We confirmed the robustness of these results (i.e., total and relative connection densities) by repeating the analyses using two different thresholds (p < 0.005 and p < 0.025) (**Table S6**). Interestingly, the analysis of the functional networks generated using Gephi software revealed a specific subnetwork involving the AOB-MOB-mPFC-OFC in the PLAY-rO group (**Fig. 7G-H**). The dHipp showed no significant correlation with any of the other selected regions. This subnetwork was not observed in CTRL-rO group. Together, these findings indicate a major reorganization of brain functional connectivity at 6 months of age in response to the childhood odorant, which is importantly relocated to the olfactory-limbic system. While the present results do not support sustained engagement of the memory system, they highlight the maintenance of a distinct functional module comprising the MOB, OFC, and mPFC, the dHipp being functionally disconnected from the network.

**Fig. 7.**
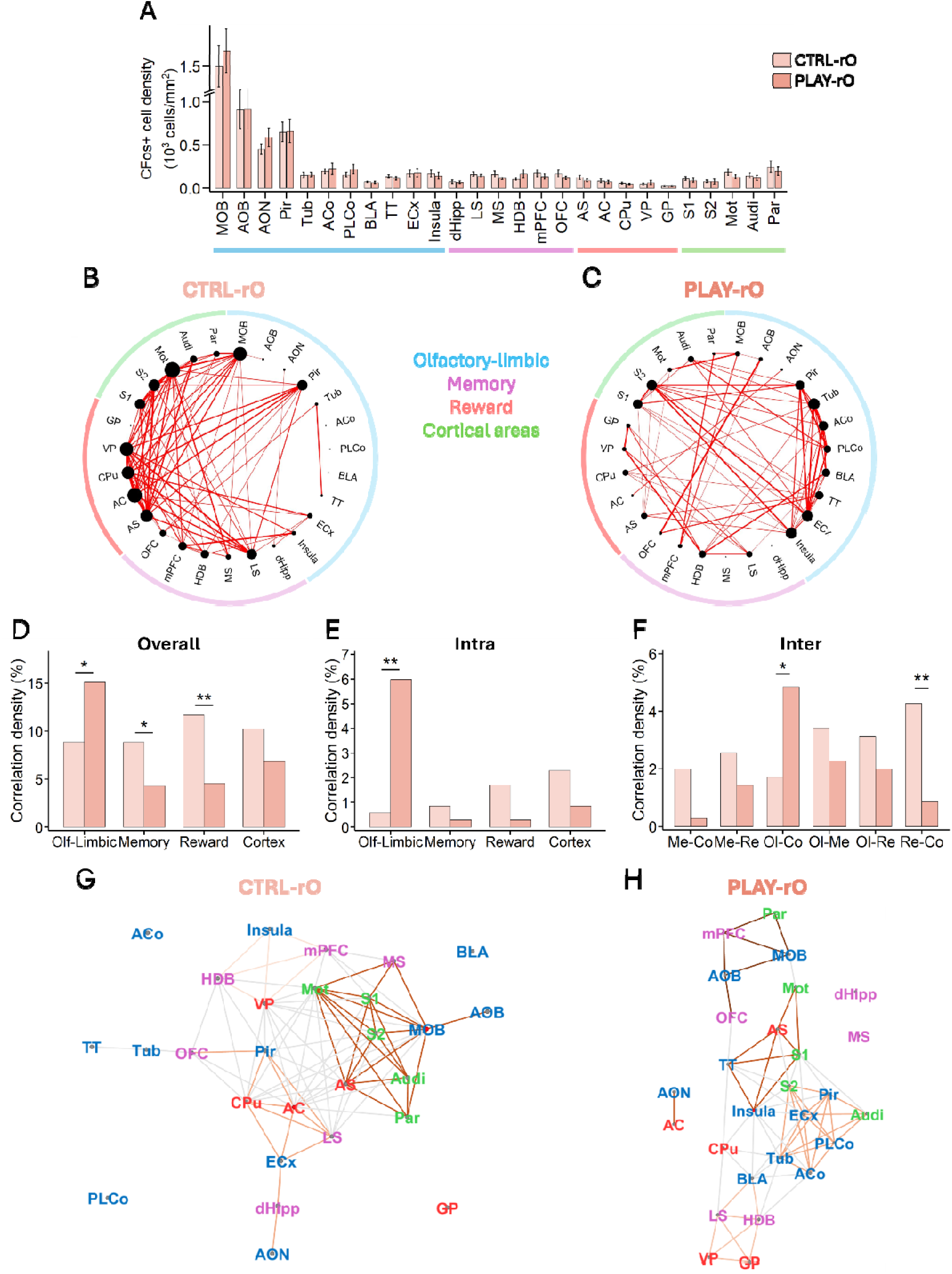
In late adulthood, the functional network recruited by the childhood odorant is reorganized around the olfactory-limbic system in late adulthood. (**A**) cFos-positive cell density is similar between PLAY-O (n=10) and CTRL-O (n=10) mice in the selected brain regions. (**B-C**) Representation of functional connectivity networks in response to the childhood odorant, constructed by considering correlations with Pearson’s p < 0.01. Each region is represented by a dot; the size of the dot is a function of the number of connections. The thickness of the line increases with the associated correlation coefficient. These networks are represented for (**B**) CTRL-rO (n=8) and (**C**) PLAY-rO (n=8) groups. (**D-F**) Total correlation density (i.e., the number of connections realized over the total possible) analysis. (**D**) The total correlation density is increased for PLAY-rO compared to CTRL-rO group in olfactory-limbic system, whereas it is decreased in memory and reward systems. (**E**) At the intra-system level, PLAY-rO group shows higher intra-olfactory-limbic connection density. (**F**) At the inter-system level, PLAY-rO group shows increased olfactory-limbic-cortex connection density compared to CTRL-rO group. (**G-H**) Representation of the Gephi networks with Louvain Communities for (**G**) CTRL-rO and (**H**) PLAY-rO groups. Text colors represent the systems to which the structures belong to. The colored lines represent the subnetworks identified with the Louvain Community approach. Statistical significance depicted as *p < 0.05, **p < 0.01. AC = Accumbens Core; ACo = Anterior Cortical Amygdala; AOB = Accessory Olfactory Bulb; AON = Anterior Olfactory Nucleus; AS = Accumbens Shell; Audi = Auditory Cortex; BLA = Basolateral Amygdala; CPu = Caudate Putamen; dHipp = dorsal Hippocampus; GP = Globus Pallidus; HDB = Horizontal Limb of the Diagonal Band of Broca; LS = Lateral Septum; MOB = Main Olfactory Bulb; Mot = Motor Cortex; mPFC = medial Prefrontal Cortex; MS = Medial Septum; OFC = Orbitofrontal Cortex; Par = Parietal Cortex; ECx = Entorhinal Cortex; Pir = Piriform Cortex; PLCo = Posterolateral Cortical Amygdala; S1 = Somatosensory Cortex 1; S2 = Somatosensory Cortex 2; Tub = Olfactory Tubercle; TT = Tenia Tecta; VP = Ventral Pallidum.

## Discussion

The primary aim of this study was to model the formation of a positive olfactory memory in juvenile mice to investigate its underlying neural basis in adulthood. Prior to the animal model, we conducted a human survey revealing that childhood olfactory memory, in addition to being emotionally positive, is most of the time encoded by a repeated event involving a pleasant odorant. To model a positive olfactory memory in mice, we repeatedly associated an odorant attractive to mice to an enriched playful environment which is reported to enhance animal well-being (Bailoo et al., 2018; Benaroya-Milshtein et al., 2004; Burgess et al., 2013). This represents an original question in three aspects. First, positive memories are rarely induced in mice, unlike fear conditioning which is more commonly used for its robustness and long-lasting effects (Frankland & Bontempi, 2005; LeDoux, 2003; Mouly & Sullivan, 2010). Second, episodic memory has been historically defined as memory for unique events (Tulving, 1972), guiding animal research to rely on single or very few repetitions to model episodic memory (Davies & Clayton, 2024). Third, our animal model was successful as it was directly inspired by a human survey and induced a positive emotional state and a long-lasting memory. Indeed, the number and frequency of ultrasonic vocalizations, which appear to be powerful measurements of a positive affective state in rodents (Granon et al., 2018; Kuwaki & Kanno, 2021; Lefebvre et al., 2020), were both increased by the exposure to the playful environment. In addition, this association during childhood led to a preference for the childhood odorant in adulthood, which can be interpreted as a positive memory of it. Importantly, we argue that this form of memory differed from familiarization since mice exposed to the childhood odorant in a playful environment (PLAY-O) showed a stronger preference for it than mice exposed to the childhood odorant without the playful environment (CTRL-O).

The secondary objective of this study was to examine how olfactory memory evolves with ageing. Although little was previously known about the dynamics of memory recall across the lifespan, our human survey revealed that the memory-associated odorant is re-experienced throughout life by the large majority (82%) of participants. In line with this finding, the mouse model demonstrates that the childhood olfactory memory is forgotten in later adulthood unless periodic re-exposures to the childhood odorant allows the maintenance of the preference.

At the neuronal level, while adult-born GCs are well known to support flexible olfactory behaviors in adulthood (Forest, Chalençon, et al., 2019; Grelat et al., 2018; Mouret et al., 2008; Sultan et al., 2011), neonatal-born GCs have been primarily associated with innate olfactory responses (Imayoshi et al., 2008; Lemasson, 2005; Muthusamy et al., 2017; Sakamoto et al., 2014; Tsuboi, 2024). Whether these early-born neurons also contribute to odor memorization during childhood, however, has remained unclear. Our findings reveal that P1-born GCs are able to sustain such olfactory memory originating from childhood, leading to a long-lasting memory into adulthood. Evidence for this comes from their higher cFos expression observed in adult, in response to the childhood odorant compared to controls (CTRL-O) and the abolition of the odorant preference upon optogenetic silencing of these neurons. Although viral transduction may extend beyond the exact injection time (i.e., P1), we argue that in our experimental conditions, most of the transduced GCs present in the adult OB originate from the early postnatal period, as this stage is concerned by a proliferation threefold larger than in adulthood and, producing GCs with very low cell death if any (Imayoshi et al., 2008; Lemasson, 2005; Platel et al., 2019; Sakamoto et al., 2014). Interestingly, this long-lasting survival may explain their ability to support a memory lasting up to adulthood.

However, in later adulthood, this memory is no longer accompanied by a preferential recruitment of P1-born GCs of the OB. Other, later-born, cohorts of GCs in the OB may have been recruited, possibly via destabilization-reconsolidation processes induced by olfactory exposures over life, or the memory engram may have been transferred to higher-order brain regions through systems consolidation, as observed for hippocampal-dependent memories (Kupke & Oliveira, 2025). In this context, analyzing brain functional connectivity enables a better comprehension of how these higher-order brain structures communicate in order to support the childhood olfactory memory throughout life.

Olfactory perception engages an extensive network downstream of the OB (Royet et al., 2003; Savic, 2005; Zald & Pardo, 2000). In young adults, the analysis of the functional connectivity between these systems revealed that the perception of the childhood odorant leads to an increased connectivity in the memory system, notably structures typically involved in autobiographical memory – the dorsal hippocampus, the prefrontal cortex – and olfactory memory – the orbitofrontal cortex and the olfactory bulb – both in humans and rodents (Reagh & Ranganath, 2018; Saive et al., 2014; Sekeres et al., 2018). Importantly, the orbitofrontal cortex is involved in processing associative olfactory learning memory in mice and odor-evoked memory in humans (Auguste et al., 2023; Masaoka et al., 2021; Matsunaga et al., 2013; Watanabe et al., 2018). In addition, the orbitofrontal cortex is reported to participate in coding the rewarding properties of sensory stimuli, being part of the mesocortical dopamine pathway (Rolls et al., 2020). More broadly, we identified an enhanced connectivity between the memory and reward systems, which is in line with the fact that odors associated with early life experiences acquire a rewarding value that promotes behaviors essential for survival (Moriceau & Sullivan, 2004). Indeed, these odors automatically orient the young animal’s behavior toward sources of safety and resources, which constitutes a fundamental adaptive mechanism.

Five months after memory encoding, exposure to the childhood odorant revealed a major functional reorganization of the brain with an enhanced connectivity in the olfactory-limbic system to the detriment of the reward system. With age, these early-life odors may progressively acquire a more sensory, and emotionally meaningful value rather than eliciting strong motivational responses. This is consistent with studies showing that odor-evoked memories often become increasingly nostalgic over time (Herz, 2016; Herz & Schooler, 2002). At the neurobiological level, this disengagement of the reward system could be driven by the prediction error due to the re-exposure to the odorant without the rewarding environment thought life leading to reduction of rewarding properties of the childhood odorant (Schultz, 2016).

The present findings further suggest that, with time, olfactory memory may shift from a detailed, episodic form toward a more perceptual representation. Importantly, while the dorsal hippocampus was still functionally recruited one month after learning, consistent with previous studies (Auguste et al., 2023; Veyrac et al., 2015), it was functionally disengaged five months later, while the typical cluster of long-term memory, involving the medial prefrontal cortex and orbitofrontal cortex, was still interconnected along with the accessory and main OB. These findings challenge the Multiple Trace Theory, which posits that the dorsal hippocampus is constantly involved in episodic memory recall (Nadel & Moscovitch, 1997; Sekeres et al., 2018). Interestingly, repeated life events may be located in a continuum between episodic and semantic memory (Addis & Szpunar, 2024). Thus, one could ask whether the repetition of the initial episode would enable the extraction of regularities from the environment, leading to a more general and abstract representation of the event. Given the role of the hippocampus in the recall of detailed episodic memories, it could be specifically disengaged from this type of memory. Further research is needed to clarify under which conditions this disengagement occurs, whether it is solely driven by the temporal dimension, or by the interplay between time and the conditions under which encoding occurs.

Taken together, these findings shed light on the ability of neonatal neurogenesis in the olfactory bulb to code a lifelong positive childhood olfactory memory, as well as on how brain regions and systems interact to support its storage and retrieval throughout life.

## Materials and Methods

### Online survey

652 participants were recruited to complete an online questionnaire. Two exclusion criteria were used for the study: (1) being outside the 18-90 age range and (2) reporting a history of olfactory disorder during childhood, which is the temporal focus of memories examined. Based on these criteria, 5 participants were excluded for reporting history of olfactory disorder during childhood, resulting in a final sample of 647 participants (174 men and 473 women), aged 18-85 years (M = 45.4, SD = 17.2). The participants were then instructed to recall and describe their earliest olfactory memory. The participants were recruited through various channels including research volunteers mailing lists, CNRS and local weekly newspapers. The online questionnaire was hosted in LimeSurvey. Participants were instructed to recall an odor that marked their childhood, going back in time as far as possible. After retrieving an olfactory autobiographical memory, they were asked to rate this memory on several self-reported measurements, that were the following: (1) The episode that triggered the memory was: (Unique / Repeated, between 2 and 5 times / Repeated, more than 5 times) (2) What is the main smell associated with this memory? (open-ended) (3) During childhood, how pleasant did you find this smell? (1 = very unpleasant, 9 = very pleasant) (4) Since then, how often have you had the opportunity to perceive this smell? (Once a week / Once a month / Once a year / Less than once a year / Never). The study was approved by the Ethical Review Committee of Inserm (n°23-1002, April 04, 2023). All the data collected were anonymized.

### Mice

A total of 272 inbred C57BL/6J male and female mice were used in this study. Mice were maintained on a 12-hour light/dark cycle at a constant temperature of 22°C, with food and water ad libitum. All procedures followed the European Community Council Directive of 22 September 2010 (2010/63/UE) and were approved by the National Ethics Committee (#46215_2023120616279607_v2). All efforts were made to minimize animal suffering.

### Experimental design

The olfactory memory formation involves associating an odorant with a positive context during early life stages. At postnatal day 21 (P21), groups of 4 or 5 mice were randomly assigned to either PLAY-O or CTRL-O conditions. At P23, PLAY-O mice were placed in a large, enriched cages designed to encourage exploration and social interactions; this environment included a wheel, tunnels, ladders, shelters and cornflakes. In CTRL-O group, mice were placed in standard cages. This was done during 2h without odorant presentation to avoid potential negative association between odorant and context due to environmental neophobia. At P24, mice were exposed to an odorant (either (+)-limonene, β-citronellol or camphor) in their regular housing for 1 h to prevent olfactory neophobia in later sessions. From P26 to P33, mice were placed in their designated experimental cage (either enriched for the PLAY-O group or standard for the CTRL-O group) for two-hour sessions every other day. During each session, the same odorant was consistently presented using a polypropylene swab impregnated with 5mL of the odorant (4 Pa, diluted in mineral oil) placed in a petri dish on the top of the cage (three petri dishes for PLAY-O condition and one for CTRL-O condition, accounting for differences in experimental cage size). As a control experiment, two other groups of mice were set up. They followed the same procedure as before, except that no odorant was introduced into the cage (PLAY-NO and CTRL-NO groups). These four experimental groups were behaviorally tested from P65 to P80. Animals were randomly assigned to each group.

### Ultrasonic vocalization recording

To assess the affective state of mice during olfactory memory formation in the experimental cage, ultrasonic vocalizations were recorded from PLAY-O and CTRL-O groups, including the cohorts of animals subsequently tested at 2 and 6 months of age. To do so, a CM16/CMPA condenser ultrasound microphone (Avisoft Biacoustics) was positioned above the cages at a sufficient height to capture sounds from at least half of the PLAY cages and the entire CTRL cages. It was connected to a personal computer via an UltraSoundGate 116Hb (Avisoft Bioacoustics), where data were recorded at a sampling frequency of 250 kHz, a buffer of 0.032s, 258 points of FFT-length, in 16-bit format. The data were then processed using fast Fourier transform in SASLab Pro software (Avisoft Biacoustics), with the following parameters: 512 FFT-length, 100% frame, Hamming window, 50%-time window overlap, resulting in a resolution of 488 Hz and 0.512 ms. Ultrasounds were manually identified, and the number, mean frequency and duration of each event were measured.

### Behavioral testing

To assess the memory of the childhood odorant in adults (2-month-old), the animals were tested in two behavioral olfactory tasks for their childhood odorant as well as another, unknown one (either (+)-limonene, β-citronellol or camphor depending on the mice). PLAY-NO and CTRL-NO groups have thus been tested for two unknown odorants, and the results were averaged.

#### Exploration test

The exploration test was used to evaluate the odorant attractiveness in a single trial. The apparatus consisted of a 40×40 cm board with an odorized central hole (Chalençon et al., 2024; Kermen et al., 2016; Mandairon et al., 2009; Midroit et al., 2021). A polypropylene swab impregnated with 60 μL of the odorant (1 Pa, diluted in mineral oil) was then placed at the bottom of a pot covered with bedding. The exploration of the hole during a 2-min trial was manually recorded.

#### Habituation test

The habituation test was used to assess the loss of interest for the odorant across repeated exposures, with the hypothesis that an increased preference would counteract the typical habituation process. The test consisted of a presentation of mineral oil, followed by four presentations of the same odorant. Each odorant presentation lasted 50 seconds and was separated by a 15-minute interval. The odorant, diluted in mineral oil (1 Pa), was presented in a tea ball. Investigation time was defined as active sniffing within 1 cm of the tea ball. The habituation slope across the four odorant presentations was then calculated.

All the behavioral analyses were done blind with regard to the experimental group.

#### Preference index

A preference index, reflecting the overall preference for the odorant, was calculated by combining the results of the two behavioral tests, using the following formula: [(x_ET_-µ_ET_)/ơ_ET_ + (x_HAB_-µ_HAB_)/ơ_HAB_] + 5 (x = animal value for a given test; the constant +5 was added to ensure all values remain positive).

#### Data Analysis

Data analyses were performed using RStudio software. Normality was assessed with Lilliefors test. Depending on the hypothesis one-tailed or two-tailed t-tests, or Wilcoxon tests, with Welch correction when needed, were then performed to compare the olfactory preference between groups.

### P1-born granule cells assessment

#### CldU injections

Mice were injected with 5-chloro-2’-deoxyuridine (CldU; Sigma Aldrich C6891) (total 50 mg/kg in saline, 3x at 2h intervals, ip) in order to specifically label a cohort of neurons born at P1 (Forest, Moreno, et al., 2019).

#### Odor stimulation and sacrifice

To investigate the neuronal activity in response to the childhood odorant, mice were exposed to a tea ball containing the childhood odorant (80 µL at 1 Pa) for 1 h. PLAY-NO and CTRL-NO groups were exposed to an unknown odorant. Then, 1 h after the odorant stimulation, mice were anesthetized with ketamine (100 mg/kg) and xylazine (10 mg/kg), then sacrificed using euthasol (50 mg/kg), followed by an intracardiac perfusion of fixative (50 mL of paraformaldehyde 4% in PBS, pH 7.4). Brains were then dissected, cryoprotected in sucrose solution (20% in PBS, 7 days), frozen rapidly (-48°C) and stored at -20°C. The brains were sectioned into 14 µm thick slices using a cryostat (Micron NX50) for cellular analysis.

#### CldU/cFos experiment

The protocol has been described previously (Forest, Moreno, et al., 2019). OB sections were incubated overnight with rat anti-CldU (1:100, Abcam, ab6326) and rabbit anti-cFos (1:2000, Abcam, 214672) antibodies at 4°C. Appropriate secondary antibodies were used (goat anti-rat Alexa 488 (1:200, Sigma Aldrich, SAB4600046), goat anti-rabbit Alexa 633 (1:200, Invitrogen, A21070)). The activity of P1-born granule cells was assessed by quantifying the percentage of CldU/cFos-positive cells in the dorsal and ventral granule cell layers of the left hemisphere (6–8 sections per animal), using a Zeiss microscope equipped with an Apotome system. On average, 98 CldU-positive cells were counted per animal. The density of CldU-positive cells was also assessed in the entire granule cell layer of the left and right hemisphere (3-8 sections per animal). The comparisons between groups were computed using t-tests or Wilcoxon tests depending on the data normality. All cells were counted blind to the experimental group.

### Optogenetic experiment

#### Surgery

To inhibit the neuronal activity of P1-born granule cells, mice were anesthetized on ice at P1, and 150 nL of pLenti-hSyn-eNpHR3.0-EYFP lentivirus (9.22 x 106 IU/mL, Addgene; Halo group) or 300 nL of control pLenti-hSyn-EYFP lentivirus (1.1 x 106 IU/mL, generated by the NGFO platform of the Lyon Neuroscience Research Center; Cont group) were injected bilaterally in the lateral ventricles, with a Hamilton syringe connected to a programmable syringe controller, at a rate of 150 nL/min (coordinates to bregma: AP +1 mm, ML +/- 1 mm, DV -2.3 mm) (Forest, Chalençon, et al., 2019). This enables the integration of the virus in proliferating cells located in the subventricular zone. From P23 to P33, all mice (Halo and Cont groups) were exposed to the PLAY-O condition as described previously. At P40, mice were injected with ketamine (100 mg/kg) and xylazine (10 mg/kg) for anesthesia and bilateral optic fibers (200 nm core diameter, 0.22 NA, Doric Lenses) were implanted in the OB with the following coordinates to bregma: AP +4.6 mm, ML +/-0.75 mm, DV -2 mm.

#### Behavior

All mice performed the two previously described behavioral tests during which they received light stimulation (crystal laser, 561 nm, 10-15 mW, continuous stimulation) when they explored the odorant (Forest, Chalençon, et al., 2019).

#### Control of light triggered inhibition

Mice were sacrificed as previously described with the exception that mice were exposed to the childhood odorant with light stimulation (0.5 s ON, 2 s OFF, pattern repeated over 30min), 1 h before sacrifice. After brain sectioning, immunochemistry was performed in OB slices with GFP/EYFP (1:750, Anaspec Tebu, 55423), cFos (1:2000, Abcam, 214672) antibodies, coupled with appropriate secondary antibodies (goat anti-chicken Alexa 488 (Invitrogen, A11039), goat anti-rabbit Alexa 546 (Invitrogen, A11010)). The density of EYFP-positive cells in the granule cell layer as well as the percentage of EYFP cells co-expressing cFos were analyzed under the optic fiber (two acquisitions on both hemispheres of 2 sections per animal). On average, 166 EYFP-positive cells were counted per animal. Verification of GFP labeling led to the exclusion of three mice (n□=□3, PLAY-O-Cont) from further analysis due to failed viral injection.

### Brain activation patterns and functional connectivity analysis

#### cFos immunochemistry

To investigate the neural networks activated in response to the childhood odorant, cFos immunochemistry was performed on brain slices (AP coordinates ranging from +6.05 to -2.59) using cFos antibody (1:2000, Abcam, 214672) followed by a biotinylated anti-rabbit secondary antibody (1/200, Vector Laboratories, BA-1000), then revealed through an avidin-biotin-peroxidase complex (ABC Elite Kit 1/100, Vector Laboratories, PK-6100) and amplified with 3,3-Diaminobenzidine (DAB 0.05%, Sigma-Aldrich, D5905). The slices were automatically scanned at the CIQLE platform (Axioscan 7).

#### cFos cell detection

cFos cell detection was realized using the open-source software QuPath. For each section, the contour of the left hemisphere and the interhemispheric line were manually drawn. Cell detection within the left hemisphere contour was carried out using the following parameters: Requested Pixel Size: 0.5, Background Radius: 8.0, Median filter radius: 1.0, Sigma: 0.7 (for the olfactory bulb, because of high cell density) or 2 (for the rest of the brain), Minimum area: 10.0, Maximum area: 200.0, Cell expansion: 0.01. The threshold was calculated based on the sections’ mean intensity. A classifier was trained to distinguish cFos-positive cells from artifacts and applied to all detected cells.

#### Atlas registration

The coordinates of each labeled cell and the contour point were exported into a software developed by the team allowing high accuracy matching of experimental brain sections with a reference brain atlas (Midroit et al., 2018; Terrier et al., 2024). This method allows precise, automatic assignment of the labeled cells to a brain structure and enables between-group comparisons after re-fitting in a common anatomical space. A total of 27 regions were selected, grouped into 4 functional systems named “olfactory-limbic”, “memory”, “reward” and “cortical areas”. The “olfactory-limbic” system was composed of 11 structures: main olfactory bulb (MOB, corresponding to the granule cell layer), accessory olfactory bulb (AOB), anterior olfactory nucleus (AON), piriform cortex (Pir), olfactory tubercle (Tub), cortical amygdala (ACo), postero-lateral cortical amygdala (PLCo), basolateral amygdala (BLA), tenia tecta (TT), entorhinal cortex (ECx) and insular cortex (Insula). The “memory” system was composed of 6 structures of interest: dorsal hippocampus (dHipp: comprising the dorsal CA1, dorsal CA3 and dentate gyrus), lateral (LS) and medial septum (MS), horizontal limb of the diagonal band of Broca (HDB), medial prefrontal cortex (mPFC : mean of the frontal and the cingulate cortex) and orbitofrontal cortex (OFC : mean of the medial and lateral orbitofrontal cortex). The “reward” system was composed of 5 structures of interest: nucleus accumbens shell (AS) and core (AC), caudate putamen (CPu), ventral pallidum (VP) and globus pallidus (GP). Finally, the “cortical areas” were composed of 5 structures of interest: somatosensory cortices 1 (S1) and 2 (S2), motor cortex (Mot), auditory cortex (Audi) and parietal cortex (Par).

#### cFos cell density analysis

For each region of interest, cFos-positive cell density was calculated by the mean of cFos-positive cell density for each slice containing the region of interest. cFos-positive cell for each was then compared between the groups using multiple Wilcoxon tests.

#### Functional connectivity analysis

Functional connectivity analysis was conducted following established method (Terrier et al., 2024; Wheeler et al., 2013). Given that correlation values are sensitive to outliers, we re-examined the acquisitions and applied a new, manual, detection threshold to animals whose mean cFos-positive cell density fell outside the μ ± 2 sd range of the overall population density (for groups tested at 2-month-old: 3 animals in PLAY-O group, 1 animal in CTRL-O group and 1 animal in PLAY-NO group). For each experimental group, the correlation coefficients of cFos-positive cell densities between each pair of regions of interest were calculated across animals, using Pearson tests (a total of 351 pairs). The coefficients were filtered to retain only significant connections based on a threshold of p < 0.01. Non-filtered correlation matrices of cFos-positive cells densities were then generated, as well as the filtered and non-filtered subtracted matrices, to visualize the group-specific connections and the differences in correlation strength, respectively. The resulting functional connectivity networks were generated, where nodes represent degrees (i.e., the number of connections), and lines thickness reflect the correlation coefficients. The total connection density, corresponding to the number of significant connections over the total number of connections possible, as well as the relative connection density, corresponding to the number of connections normalized by total connections observed in each group, were calculated for each system (olfactory, mnesic and reward systems and cortical areas). To assess the robustness of the results, the same analysis was conducted with two other p-value thresholds (p < 0.025, p < 0.005). Another representation was computed with Gephi software, allowing the representation of Louvain Communities. The Louvain algorithm operates in two iterative phases: initially, each node is assigned to its own community, then it is moved to the neighboring community that maximizes the modularity gain (i.e., assess how well a network can be partitioned into communities of modules). A new graph is then created by aggregating the detected communities into single nodes, and the process repeats until no further increase in modularity is detected. The communities identified by the Louvain algorithm reflect a modular organization of the brain, where groups of brain structures preferentially collaborate, suggesting local functional specialization within a globally integrated system.

### Assessment of long-term retention of the childhood olfactory memory

The memory of the childhood odorant was also assessed in four additional cohorts of animals, at 6 months of age. In the first two groups, animals were exposed to either the PLAY environment (PLAY-O) or a standard cage (CTRL-O) during childhood, paired with an odorant as previously described. The other two groups received the same treatment except they were re-exposed to the childhood odorant for 2 hours every 3 weeks in their regular housing between P65 and P80 (7 re-exposures in total, PLAY-rO and CTRL-rO). As previously described, olfactory preference was assessed in response to the childhood and an unknown odorant. The activity of P1-born GCs has also been assessed, as well as the functional connectivity analysis.

## Acknowledgments

The author(s) declare that financial support was received for the research, authorship, and/or publication of this article. This work was supported by Observatoire B2V des Mémoires (doctoral fellowship to J.D.), CNRS, ANR-23-CZE17-0065-01, INSERM, Claude Bernard Lyon 1 University, IUF. Slice acquisitions were performed using microscopes funded by the Auvergne-Rhône-Alpes Region, as well as on the CIQLE platform. The manuscript was edited for language and clarity with the assistance of ChatGPT.

## Author Contributions

Conceptualization: A.D., J.D. and N.M.

Investigation: A.A, J.D. and T.B.

Software: M.T.

Visualization: J.D.

Formal analysis: J.D. and M.T.

Writing – original draft: J.D.

Writing – review & editing: A.D., J.D. and N.M.

## Competing Interest Statement

The authors declare that the research was conducted in the absence of any commercial or financial relationships that could be construed as a potential conflict of interest.

## Notes

### Competing Interest Statement

The authors have declared no competing interest.

### Summary of Updates

The title as well as the abstract have been revised.

