## Supplementary material for "Positive olfactory childhood memory is rooted in the olfactory bulb and triggers large scale changes beyond the olfactory system": Fig. S1

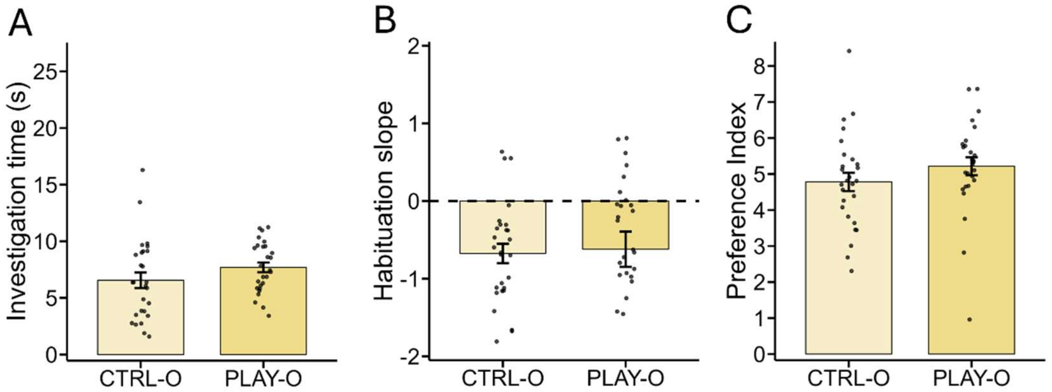


**Fig. S1. Behavioral responses to an unknown odorant in 2-month-old mice (PLAY-O and CTRL-O groups).** No difference is found between PLAY-O and CTRL-O mice regarding (**A**) the investigation time in the exploration test, (**B**) the habituation slope in the habituation test, nor (**C**) the resulting preference index. Data are represented as data points and mean ± SEM.
