## Supplementary material for "Positive olfactory childhood memory is rooted in the olfactory bulb and triggers large scale changes beyond the olfactory system": Fig. S2

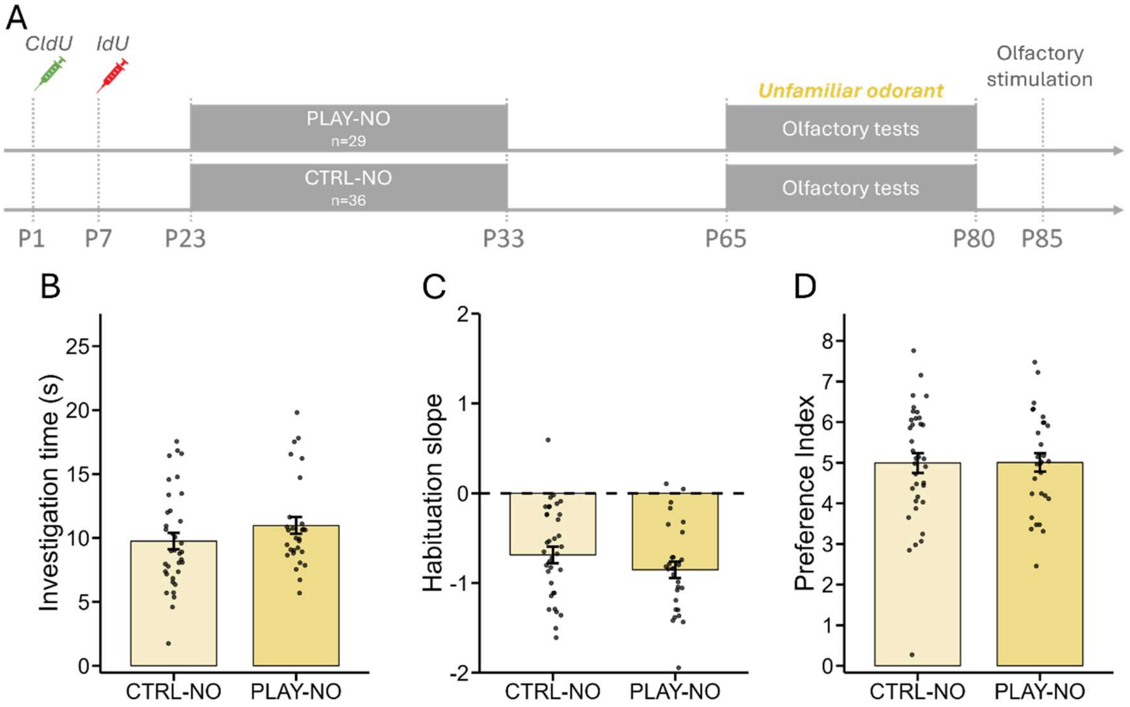


**Fig. S2. Behavioral responses to an unknown odorant in 2-month-old mice (PLAY-NO and CTRL-NO groups).** (**A**) Timeline of the experiment. PLAY-NO (n=29) and CTRL-NO (n=36) mice show similar (**B**) investigation time in the exploration test, (**C**) habituation slope in the habituation test as well as (**D**) resulting preference index. Data are represented as data points and mean ± SEM.
