## Supplementary material for "Positive olfactory childhood memory is rooted in the olfactory bulb and triggers large scale changes beyond the olfactory system": Fig. S3

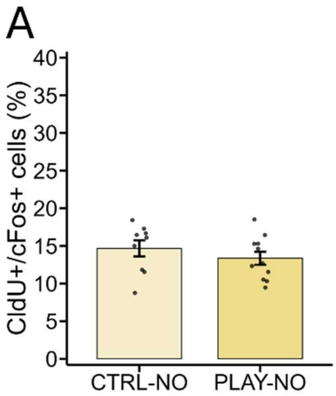


**Fig. S3. Cellular analyses in 2-month-old mice.** (**A**) The percentage of CldU-positive cells responding to an unknown odorant are similar between PLAY-NO (n=11) and CTRL-NO (n=9) mice. Data are represented as data points and mean ± SEM.
