## Supplementary material for "Positive olfactory childhood memory is rooted in the olfactory bulb and triggers large scale changes beyond the olfactory system": Fig. S4

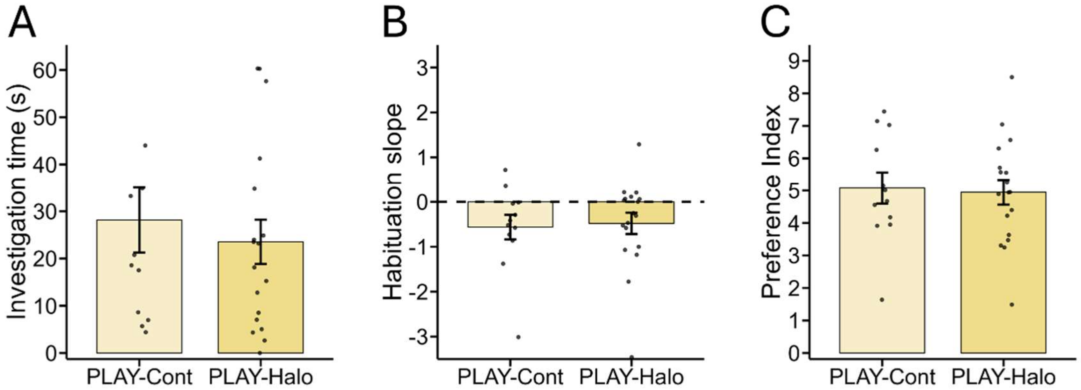


**Fig. S4. Behavioral responses to an unknown odorant in the optogenetics experiment.** The light-induced inhibition of P1-born GCs (PLAY-Halo group, n=18) does not affect (**A**) the investigation time, (**B**) the habituation slope, nor (**C**) the preference index in response to an unknown odorant, compared to the PLAY-Cont group (n=12). Data are represented as data points and mean ± SEM.
