## Supplementary material for "Positive olfactory childhood memory is rooted in the olfactory bulb and triggers large scale changes beyond the olfactory system": Fig. S5

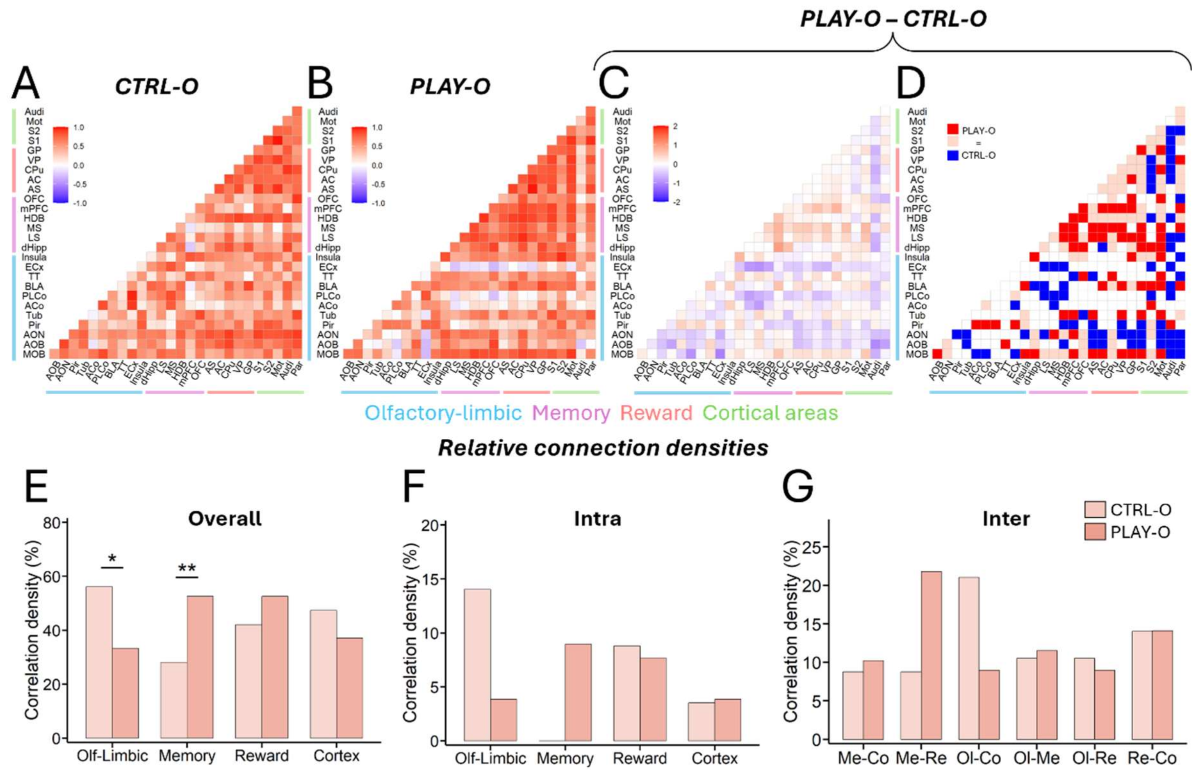


**Fig. S5. Functional connectivity analysis in 2-month-old mice (PLAY-O and CTRL-O groups).** (**A-B**) Correlation matrices without p-value thresholding for (**A**) CTRL-O and (**B**) PLAY-O groups. Matrix color represents the correlation coefficients between pairs of regions, ranging from -1 to 1. (**C-D**) Group comparison matrices (PLAY-O – CTRL-O) on (**C**) non-thresholded matrices to visualize differences in correlation coefficients (scale: -2 to +2; positive values indicate higher correlation coefficients in the PLAY-O group) and on (**D**) thresholded ($p < 0.01$) matrices to visualize group-specific correlations (red = specific to PLAY-O, blue = specific to CTRL-O, pink = shared correlations). (**E-G**) Relative correlation density (i.e., the number of connections normalized by total connections observed in each group) analysis. (**E**) The relative correlation density is decreased in the olfactory-limbic system and increased in the memory system for PLAY-O compared to CTRL-O group. No significant group differences are observed at the (**F**) intra-system nor (**G**) inter-system level. Statistical significance depicted as $^*p < 0.05$, $^{**}p < 0.01$. *AC = Accumbens Core; ACo = Anterior Cortical Amygdala; AOB = Accessory Olfactory Bulb; AON = Anterior Olfactory Nucleus; AS = Accumbens Shell; Audi = Auditory Cortex; BLA = Basolateral Amygdala; CPu = Caudate Putamen; dHipp = dorsal Hippocampus; GP = Globus Pallidus; HDB = Horizontal Limb of the Diagonal Band of Broca; LS = Lateral Septum; MOB = Main Olfactory Bulb; Mot = Motor Cortex; mPFC = medial Prefrontal Cortex; MS = Medial Septum; OFC = Orbitofrontal Cortex; Par = Parietal Cortex; ECx = Entorhinal Cortex; Pir = Piriform Cortex; PLCo = Posterolateral Cortical Amygdala; S1 = Somatosensory Cortex 1; S2 = Somatosensory Cortex 2; Tub = Olfactory Tubercle; TT = Tenia Tecta; VP = Ventral Pallidum.*
