## Supplementary material for "Positive olfactory childhood memory is rooted in the olfactory bulb and triggers large scale changes beyond the olfactory system": Fig. S6

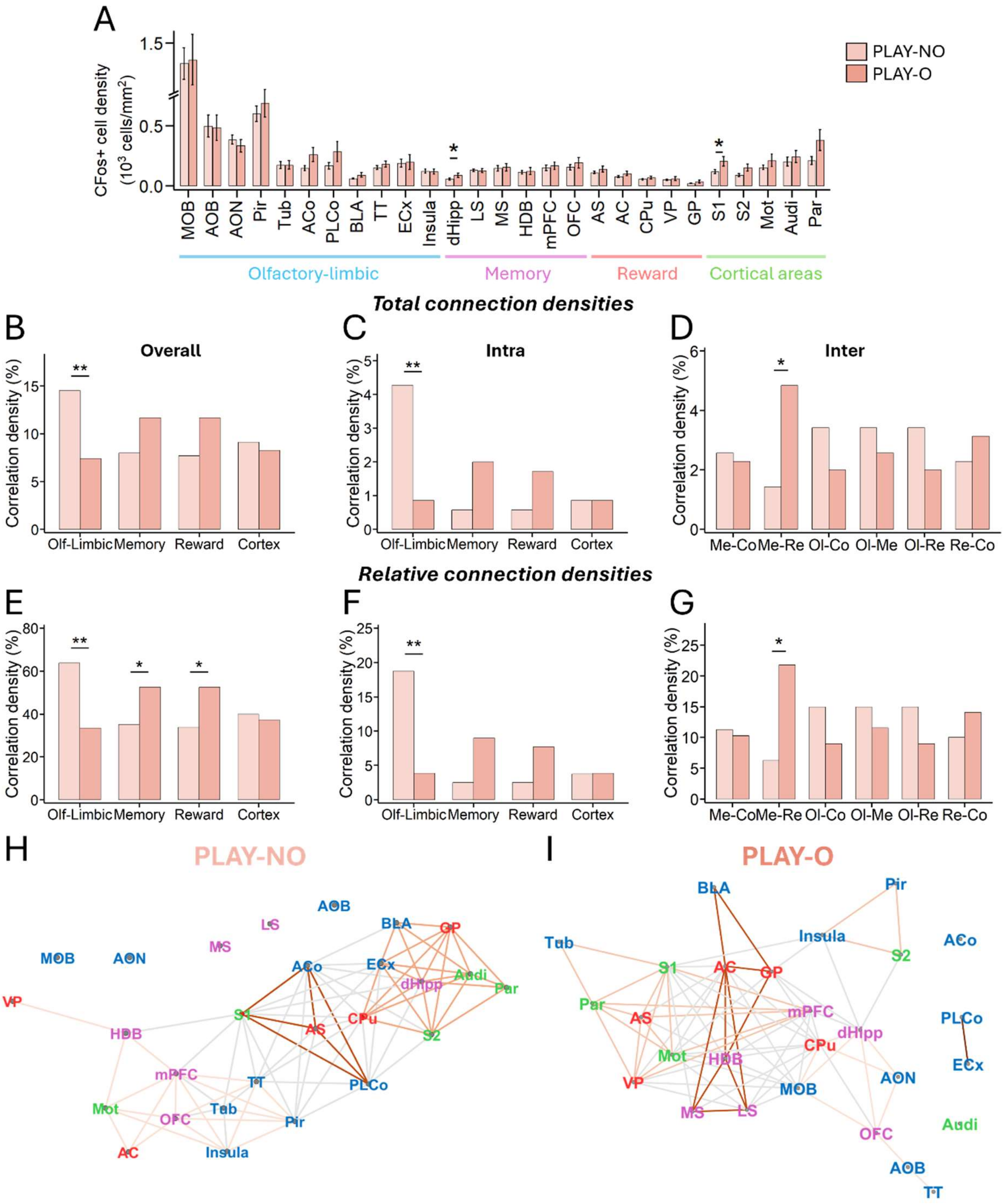


**Fig. S6. Functional connectivity analysis in 2-month-old mice (PLAY-O and PLAY-NO groups).** (**A**) cFos-positive cell density is higher in the dHipp and S1 for PLAY-O (n=10) compared to PLAY-NO (n=8) mice. (**B-D**) Total correlation density analysis (i.e., the number of connections realized over the total possible) analysis. (**B**) The correlation density is decreased in the olfactory-limbic system for the PLAY-O compared to the PLAY-NO group, reflected by (**C**) a decrease in the intra olfactory-limbic system. (**D**) The PLAY-O group shows increased inter memory-reward correlation density compared to the PLAY-NO group. (**E-G**) Relative correlation density (i.e., the number of connections normalized by total connections observed in each group) analysis. (**E**) The relative correlation density is decreased in the olfactory-limbic system and increased in the memory and reward system for the PLAY-O compared to the PLAY-NO group. These differences are reflected by (**F**) a lower correlation density in the intra-olfactory-limbic system and (**G**) a higher memory-reward correlation density for the PLAY-O compared to the PLAY-NO group. (**H-I**) Representation of the Gephi networks with Louvain Communities for (**H**) PLAY-NO and (**I**) PLAY-O groups. Text colors represent the systems to which the structures belong. The colored lines represent the subnetworks identified with the Louvain Community approach. Statistical significance depicted as *p < 0.05, **p < 0.01. *AC = Accumbens Core; ACo = Anterior Cortical Amygdala; AOB = Accessory Olfactory Bulb; AON = Anterior Olfactory Nucleus; AS = Accumbens Shell; Audi = Auditory Cortex; BLA = Basolateral Amygdala; CPu = Caudate Putamen; dHipp = dorsal Hippocampus; GP = Globus Pallidus; HDB = Horizontal Limb of the Diagonal Band of Broca; LS = Lateral Septum; MOB = Main Olfactory Bulb; Mot = Motor Cortex; mPFC = medial Prefrontal Cortex; MS = Medial Septum; OFC = Orbitofrontal Cortex; Par = Parietal Cortex; ECx = Entorhinal Cortex; Pir = Piriform Cortex; PLCo = Posterolateral Cortical Amygdala; S1 = Somatosensory Cortex 1; S2 = Somatosensory Cortex 2; Tub = Olfactory Tubercle; TT = Tenia Tecta; VP = Ventral Pallidum.*
