## Supplementary material for "Positive olfactory childhood memory is rooted in the olfactory bulb and triggers large scale changes beyond the olfactory system": Fig. S7

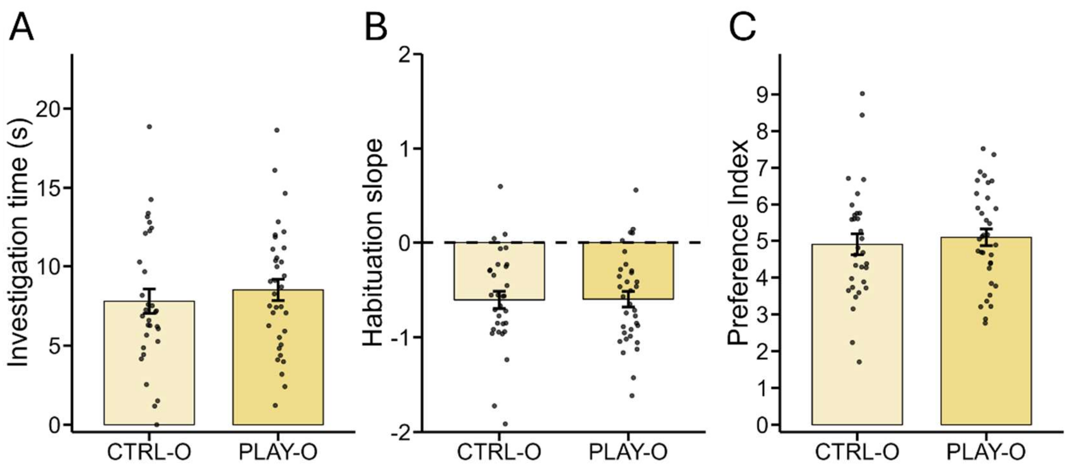


**Fig. S7. Behavioral results in 6-month-old mice.** (**A-C**) Behavioral responses to an unknown odorant. At 6 months of age, PLAY-O (n=34) and CTRL-O (n=31) groups show similar (**A**) habituation slope in the habituation test, (**B**) investigation time in the exploration test as well as (**C**) resulting preference index. Data are represented as data points and mean ± SEM.
