## Supplementary material for "Positive olfactory childhood memory is rooted in the olfactory bulb and triggers large scale changes beyond the olfactory system": Fig. S8

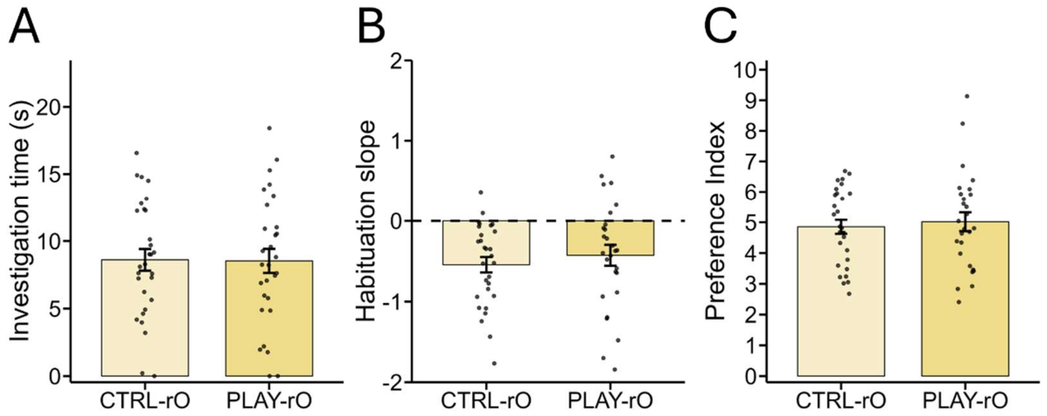


**Fig. S8. Behavioral results in 6-month-old mice (with periodic olfactory re-exposures).** (**A-C**) Behavioral responses to an unknown odorant. PLAY-rO (n=29) and CTRL-rO (n=29) groups show similar (**A**) investigation time, (**B**) habituation slope and (**C**) preference index in response to an unknown odorant. Data are represented as data points and mean ± SEM.
