## Supplementary material for "Positive olfactory childhood memory is rooted in the olfactory bulb and triggers large scale changes beyond the olfactory system": Fig. S9

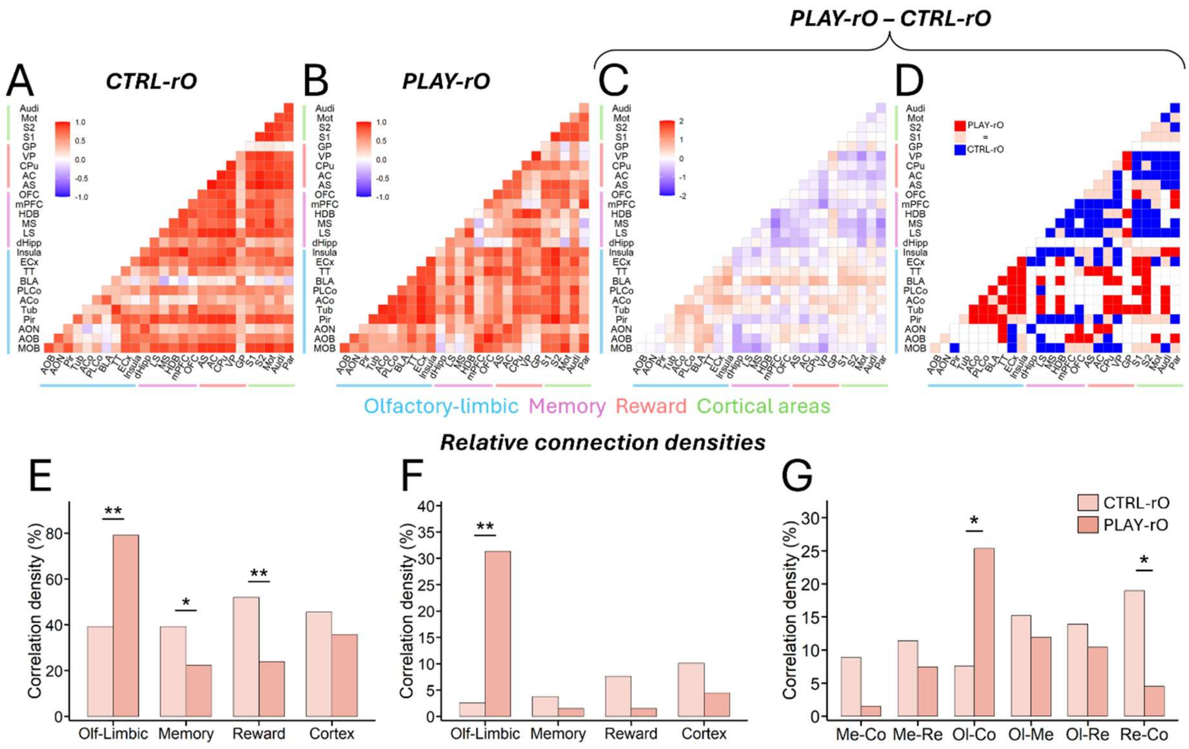


**Fig. S9. Functional connectivity analysis in 6-month-old mice (with periodic re-exposures).** (**A-B**) Correlation matrices without p-value thresholding for (**A**) CTRL-rO and (**B**) PLAY-rO groups. Matrix color represents the correlation coefficients between pairs of regions, ranging from -1 to 1. (**C-D**) Group comparison matrices (PLAY-rO – CTRL-rO) on (**C**) non-thresholded matrices to visualize differences in correlation coefficients (scale: -2 to +2; positive values indicate higher correlation coefficients in the PLAY-O group) and on (**D**) thresholded (p < 0.01) matrices to visualize group-specific correlations (red = specific to PLAY-rO, blue = specific to CTRL-rO, pink = shared correlations). (**E-G**) The relative correlation density (i.e., the number of connections normalized by total connections observed in each group) is (**E**), increased in the olfactory-limbic system and decreased in the memory and reward systems for the PLAY-rO compared to the CTRL-rO group. These differences are reflected by (**F**) a higher correlation density in the intra-olfactory-limbic system for the PLAY-rO group as well as (**G**) a higher olfactory-limbic-cortex and a lower reward-cortex correlation density for the PLAY-rO compared to the CTRL-rO group. Statistical significance depicted as *p < 0.05, **p < 0.01. *AC = Accumbens Core; ACo = Anterior Cortical Amygdala; AOB = Accessory Olfactory Bulb; AON = Anterior Olfactory Nucleus; AS = Accumbens Shell; Audi = Auditory Cortex; BLA = Basolateral Amygdala; CPu = Caudate Putamen; dHipp = dorsal Hippocampus; GP = Globus Pallidus; HDB = Horizontal Limb of the Diagonal Band of Broca; LS = Lateral Septum; MOB = Main Olfactory Bulb; Mot = Motor Cortex; mPFC = medial Prefrontal Cortex; MS = Medial Septum; OFC = Orbitofrontal Cortex; Par = Parietal Cortex; ECx = Entorhinal Cortex; Pir = Piriform Cortex; PLCo = Posterolateral Cortical Amygdala; S1 = Somatosensory Cortex 1; S2 = Somatosensory Cortex 2; Tub = Olfactory Tubercle; TT = Tenia Tecta; VP = Ventral Pallidum.*
