## Supplementary material for "Positive olfactory childhood memory is rooted in the olfactory bulb and triggers large scale changes beyond the olfactory system": Table S1

**Table S1. Comparison of cFos-positive cell densities between PLAY-O and CTRL-O groups, tested at two months.**

| Region | MOB | AOB | AON | Pir | Tub | ACo | PLCo | BLA | TT | ECx | Insula | dHipp | LS | MS |
|---|---|---|---|---|---|---|---|---|---|---|---|---|---|---|
| p-value | 0.739 | 0.190 | 0.579 | 0.739 | 0.971 | 0.218 | 0.796 | 0.247 | 0.912 | 0.315 | 0.853 | 0.353 | 0.280 | 0.481 |
| **Region** | **HDB** | **mPFC** | **OFC** | **AS** | **AC** | **CPu** | **VP** | **GP** | **S1** | **S2** | **Mot** | **Audi** | **Par** | |
| p-value | 0.436 | 0.579 | 0.684 | 0.853 | 0.912 | 0.853 | 0.315 | 0.579 | 0.796 | 0.796 | 0.579 | 0.739 | 0.315 | |
