## Supplementary material for "Positive olfactory childhood memory is rooted in the olfactory bulb and triggers large scale changes beyond the olfactory system": Table S2

**Table S2. Functional connectivity analyses between PLAY-O and CTRL-O groups, tested at two months.**

| | | p < 0.005 | | p < 0.01 | | p < 0.025 | |
|---|---|---|---|---|---|---|---|
| | **System** | **X-square** | **p-value** | **X-square** | **p-value** | **X-square** | **p-value** |
| **Total overall** | All | 2.550 | 0.1103 | 3.607 | 0.0575 | 0.235 | 0.6276 |
| | Olfactory-limbic | 0.249 | 0.6175 | 0.467 | 0.4944 | 2.016 | 0.1556 |
| | Memory | 6.313 | 0.0120 | 10.929 | 0.0009 | 5.196 | 0.0226 |
| | Reward | 3.370 | 0.0664 | 4.309 | 0.0379 | 1.444 | 0.2295 |
| | Cortex | 0.024 | 0.8766 | 0.019 | 0.8896 | 0.984 | 0.3212 |
| **Total intra** | Olfactory-limbic | 1.512 | 0.2188 | 1.476 | 0.2244 | 0.512 | 0.4742 |
| | Memory | 4.200 | 0.0404 | 5.191 | 0.0227 | 3.321 | 0.0684 |
| | Reward | 0.000 | 1.0000 | 0.000 | 1.0000 | 0.313 | 0.5758 |
| | Cortex | 0.000 | 1.0000 | 0.000 | 1.0000 | 0.000 | 1.0000 |
| **Total inter** | Memory-Cortex | 0.000 | 1.0000 | 0.313 | 0.5758 | 0.784 | 0.3760 |
| | Memory-Reward | 2.789 | 0.0949 | 5.665 | 0.0173 | 3.586 | 0.0583 |
| | Olfactory-limbic-Cortex | 0.313 | 0.5758 | 0.864 | 0.3527 | 2.812 | 0.0936 |
| | Olfactory-limbic-Memory | 0.101 | 0.7502 | 0.272 | 0.6020 | 0.040 | 0.8418 |
| | Olfactory-limbic-Reward | 0.000 | 1.0000 | 0.000 | 1.0000 | 0.047 | 0.8287 |
| | Reward-Cortex | 0.512 | 0.4742 | 0.216 | 0.6421 | 0.294 | 0.5879 |
| **Relative overall** | Olfactory-limbic | 3.453 | 0.0632 | 6.091 | 0.0136 | 4.955 | 0.0260 |
| | Memory | 3.531 | 0.0602 | 7.126 | 0.0076 | 5.715 | 0.0168 |
| | Reward | 0.828 | 0.3629 | 1.054 | 0.3045 | 1.168 | 0.2797 |
| | Cortex | 0.624 | 0.4296 | 1.020 | 0.3125 | 2.786 | 0.0951 |
| **Relative intra** | Olfactory-limbic | 2.820 | 0.0931 | 3.308 | 0.0689 | 0.784 | 0.3760 |
| | Memory | 2.991 | 0.0837 | 3.724 | 0.0536 | 3.051 | 0.0807 |
| | Reward | 0.000 | 1.0000 | 0.000 | 1.0000 | 0.203 | 0.6524 |
| | Cortex | 0.000 | 1.0000 | 0.000 | 1.0000 | 0.013 | 0.9108 |
| **Relative inter** | Memory-Cortex | 0.000 | 1.0000 | 0.000 | 1.0000 | 0.579 | 0.4466 |
| | Memory-Reward | 1.206 | 0.2720 | 3.196 | 0.0738 | 3.271 | 0.0705 |
| | Olfactory-limbic-Cortex | 1.475 | 0.2246 | 3.037 | 0.0814 | 3.873 | 0.0491 |
| | Olfactory-limbic-Memory | 0.000 | 1.0000 | 0.000 | 1.0000 | 0.151 | 0.6977 |
| | Olfactory-limbic-Reward | 0.000 | 1.0000 | 0.000 | 0.9948 | 0.004 | 0.9502 |
| | Reward-Cortex | 0.002 | 0.9630 | 0.000 | 1.0000 | 0.604 | 0.4369 |
