## Supplementary material for "Positive olfactory childhood memory is rooted in the olfactory bulb and triggers large scale changes beyond the olfactory system": Table S3

**Table S3. Comparison of cFos-positive cell densities between PLAY-O and PLAY-NO groups, tested at two months.**

| Region | MOB | AOB | AON | Pir | Tub | ACo | PLCo | BLA | TT | ECx | Insula | dHipp | LS | MS |
|---|---|---|---|---|---|---|---|---|---|---|---|---|---|---|
| p-value | 0.958 | 0.715 | 0.433 | 0.520 | 0.590 | 0.087 | 0.227 | 0.286 | 0.271 | 0.931 | 0.741 | 0.041 | 0.903 | 0.543 |

| Region | HDB | mPFC | OFC | AS | AC | CPu | VP | GP | S1 | S2 | Mot | Audi | Par |
|---|---|---|---|---|---|---|---|---|---|---|---|---|---|
| p-value | 0.958 | 0.475 | 0.520 | 0.433 | 0.303 | 0.566 | 0.958 | 0.303 | 0.045 | 0.101 | 0.433 | 0.320 | 0.063 |
