## Supplementary material for "Positive olfactory childhood memory is rooted in the olfactory bulb and triggers large scale changes beyond the olfactory system": Table S4

**Table S4. Functional connectivity analyses between PLAY-O and PLAY-NO groups, tested at two months.**

| | | p < 0.005 | | p < 0.01 | | p < 0.025 | |
|---|---|---|---|---|---|---|---|
| | System | X-square | p-value | X-square | p-value | X-square | p-value |
| Total overall | All | 0.010 | 0.9207 | 0.008 | 0.9287 | 0.752 | 0.3857 |
| | Olfactory-limbic | 8.505 | 0.0035 | 8.329 | 0.0039 | 14.208 | 0.0002 |
| | Memory | 1.735 | 0.1878 | 2.297 | 0.1297 | 1.444 | 0.2295 |
| | Reward | 2.824 | 0.0929 | 2.731 | 0.0984 | 0.565 | 0.4524 |
| | Cortex | 0.000 | 1.0000 | 0.071 | 0.7894 | 0.116 | 0.7337 |
| Total intra | Olfactory-limbic | 8.806 | 0.0030 | 6.886 | 0.0087 | 6.268 | 0.0123 |
| | Memory | 1.137 | 0.2863 | 1.799 | 0.1798 | 3.321 | 0.0684 |
| | Reward | 0.577 | 0.4476 | 1.137 | 0.2863 | 1.476 | 0.2244 |
| | Cortex | 0.000 | 1.0000 | 0.000 | 1.0000 | 0.000 | 1.0000 |
| Total inter | Memory-Cortex | 0.000 | 1.0000 | 0.000 | 1.0000 | 0.421 | 0.5163 |
| | Memory-Reward | 5.172 | 0.0230 | 5.665 | 0.0173 | 2.153 | 0.1423 |
| | Olfactory-limbic-Cortex | 0.085 | 0.7711 | 0.864 | 0.3527 | 2.812 | 0.0936 |
| | Olfactory-limbic-Memory | 0.575 | 0.4484 | 0.196 | 0.6580 | 1.599 | 0.2061 |
| | Olfactory-limbic-Reward | 0.655 | 0.4183 | 0.864 | 0.3527 | 1.211 | 0.2711 |
| | Reward-Cortex | 0.512 | 0.4742 | 0.216 | 0.6421 | 0.000 | 1.0000 |
| Relative overall | Olfactory-limbic | 13.486 | 0.0002 | 13.433 | 0.0002 | 17.450 | < 0.0001 |
| | Memory | 3.316 | 0.0686 | 4.265 | 0.0389 | 4.542 | 0.0331 |
| | Reward | 5.602 | 0.0179 | 4.961 | 0.0259 | 2.611 | 0.1061 |
| | Cortex | 0.020 | 0.8875 | 0.040 | 0.8410 | 0.000 | 1.0000 |
| Relative intra | Olfactory-limbic | 9.376 | 0.0022 | 7.277 | 0.0070 | 5.434 | 0.0197 |
| | Memory | 1.315 | 0.2514 | 1.994 | 0.1579 | 4.129 | 0.0422 |
| | Reward | 0.680 | 0.4096 | 1.267 | 0.2604 | 1.998 | 0.1576 |
| | Cortex | 0.000 | 1.0000 | 0.000 | 1.0000 | 0.025 | 0.8741 |
| Relative inter | Memory-Cortex | 0.000 | 1.0000 | 0.000 | 1.0000 | 0.862 | 0.3531 |
| | Memory-Reward | 6.194 | 0.0128 | 6.718 | 0.0095 | 3.443 | 0.0635 |
| | Olfactory-limbic-Cortex | 0.059 | 0.8076 | 0.846 | 0.3578 | 2.130 | 0.1445 |
| | Olfactory-limbic-Memory | 0.539 | 0.4629 | 0.165 | 0.6844 | 1.038 | 0.3082 |
| | Olfactory-limbic-Reward | 0.619 | 0.4314 | 0.846 | 0.3578 | 0.722 | 0.3956 |
| | Reward-Cortex | 0.712 | 0.3988 | 0.300 | 0.5837 | 0.082 | 0.7746 |
