## Supplementary material for "Positive olfactory childhood memory is rooted in the olfactory bulb and triggers large scale changes beyond the olfactory system": Table S5

**Table S5. Comparison of cFos-positive cell densities between PLAY-rO and CTRL-rO groups, tested at six months.**

| Region | MOB | AOB | AON | Pir | Tub | ACo | PLCo | BLA | TT | ECx | Insula | dHipp | LS | MS |
|---|---|---|---|---|---|---|---|---|---|---|---|---|---|---|
| p-value | 0.721 | 0.878 | 0.195 | 0.721 | 1.000 | 0.574 | 0.645 | 0.382 | 0.645 | 0.505 | 0.505 | 0.878 | 0.328 | 0.645 |
| **Region** | **HDB** | **mPFC** | **OFC** | **AS** | **AC** | **CPu** | **VP** | **GP** | **S1** | **S2** | **Mot** | **Audi** | **Par** | |
| p-value | 0.721 | 0.574 | 0.442 | 0.382 | 0.721 | 0.505 | 1.000 | 0.721 | 0.574 | 0.721 | 0.442 | 0.382 | 0.721 | |
