## Supplementary material for "Positive olfactory childhood memory is rooted in the olfactory bulb and triggers large scale changes beyond the olfactory system": Table S6

**Table S6. Functional connectivity analyses between PLAY-rO and CTRL-rO groups, tested at six months.**

| | | p < 0.005 | | p < 0.01 | | p < 0.025 | |
|---|---|---|---|---|---|---|---|
| | System | X-square | p-value | X-square | p-value | X-square | p-value |
| Total overall | All | 4.386 | 0.0362 | 1.027 | 0.3108 | 2,394 | 0,1218 |
| | Olfactory-limbic | 1.735 | 0.1878 | 5.906 | 0.0151 | 3,576 | 0,0586 |
| | Memory | 1.489 | 0.2223 | 5.208 | 0.0225 | 8,548 | 0,0035 |
| | Reward | 23.210 | < 0.0001 | 10.929 | 0.0009 | 8,935 | 0,0028 |
| | Cortex | 5.299 | 0.0213 | 2.191 | 0.1389 | 3,078 | 0,0794 |
| Total intra | Olfactory-limbic | 10.791 | 0.0010 | 14.529 | 0.0001 | 7,275 | 0,0070 |
| | Memory | 0.000 | 1.0000 | 0.251 | 0.6161 | 2,307 | 0,1288 |
| | Reward | 2.307 | 0.1288 | 2.307 | 0.1288 | 0,450 | 0,5024 |
| | Cortex | 2.307 | 0.1288 | 1.476 | 0.2244 | 0,272 | 0,6020 |
| Total inter | Memory-Cortex | 0.251 | 0.6161 | 3.158 | 0.0755 | 3,566 | 0,0590 |
| | Memory-Reward | 1.476 | 0.2244 | 0.655 | 0.4183 | 1,170 | 0,2793 |
| | Olfactory-limbic-Cortex | 0.575 | 0.4484 | 4.484 | 0.0342 | 2,643 | 0,1040 |
| | Olfactory-limbic-Memory | 0.000 | 1.0000 | 0.462 | 0.4966 | 0,662 | 0,4159 |
| | Olfactory-limbic-Reward | 5.904 | 0.0151 | 0.512 | 0.4742 | 0,040 | 0,8418 |
| | Reward-Cortex | 10.246 | 0.0014 | 6.886 | 0.0087 | 9,600 | 0,0019 |
| Relative overall | Olfactory-limbic | 17.260 | < 0.0001 | 21.977 | < 0.0001 | 18,541 | < 0.0001 |
| | Memory | 0.026 | 0.8708 | 4.022 | 0.0449 | 5,976 | 0,0145 |
| | Reward | 20.511 | < 0.0001 | 10.810 | 0.0010 | 6,341 | 0,0118 |
| | Cortex | 1.492 | 0.2218 | 1.049 | 0.3057 | 1,029 | 0,3105 |
| Relative intra | Olfactory-limbic | 21.230 | < 0.0001 | 20.557 | < 0.0001 | 12,692 | 0,0004 |
| | Memory | 0.000 | 1.0000 | 0.117 | 0.7328 | 1,652 | 0,1987 |
| | Reward | 1.019 | 0.3127 | 1.772 | 0.1832 | 0,151 | 0,6977 |
| | Cortex | 1.019 | 0.3127 | 0.949 | 0.3300 | 0,023 | 0,8806 |
| Relative inter | Memory-Cortex | 0.006 | 0.9379 | 2.511 | 0.1131 | 2,471 | 0,1160 |
| | Memory-Reward | 0.297 | 0.5858 | 0.272 | 0.6020 | 0,427 | 0,5135 |
| | Olfactory-limbic-Cortex | 3.236 | 0.0720 | 7.346 | 0.0067 | 5,437 | 0,0197 |
| | Olfactory-limbic-Memory | 0.359 | 0.5493 | 0.107 | 0.7433 | 0,117 | 0,7324 |
| | Olfactory-limbic-Reward | 3.454 | 0.0631 | 0.148 | 0.7009 | 0,000 | 1,0000 |
| | Reward-Cortex | 7.139 | 0.0075 | 5.783 | 0.0162 | 7,674 | 0,0056 |
